# An intact amber-free HIV-1 system for in-virus protein bioorthogonal click labeling that delineates envelope conformational dynamics

**DOI:** 10.1101/2023.02.28.530526

**Authors:** Yuanyun Ao, Jonathan R. Grover, Yang Han, Guohua Zhong, Wenyi Qin, Dibya Ghimire, Anzarul Haque, Rajanya Bhattacharjee, Baoshan Zhang, James Arthos, Edward A. Lemke, Peter D. Kwong, Maolin Lu

**Author notes:** Correspondence should be addressed to M.L.

## Abstract

The HIV-1 envelope (Env) glycoprotein is conformationally dynamic and mediates membrane fusion required for cell entry. Single-molecule fluorescence resonance energy transfer (smFRET) of Env using peptide tags has provided mechanistic insights into the dynamics of Env conformations. Nevertheless, using peptide tags risks potential effects on structural integrity. Here, we aim to establish minimally invasive smFRET systems of Env on the virus by combining genetic code expansion and bioorthogonal click chemistry. Amber stop-codon suppression allows site-specifically incorporating noncanonical/unnatural amino acids (ncAAs) at introduced amber sites into proteins. However, ncAA incorporation into Env (or other HIV-1 proteins) in the virus context has been challenging due to low copies of Env on virions and incomplete amber suppression in mammalian cells. Here, we developed an intact amber-free virus system that overcomes impediments from preexisting ambers in HIV-1. Using this system, we successfully incorporated dual ncAAs at amber-introduced sites into Env on intact virions. Dual-ncAA incorporated Env retained similar neutralization sensitivities to neutralizing antibodies as wildtype. smFRET of click-labeled Env on intact amber-free virions recapitulated conformational profiles of Env. The amber-free HIV-1 infectious system also permits in-virus protein bioorthogonal labeling, compatible with various advanced microscopic studies of virus entry, trafficking, and egress in living cells.

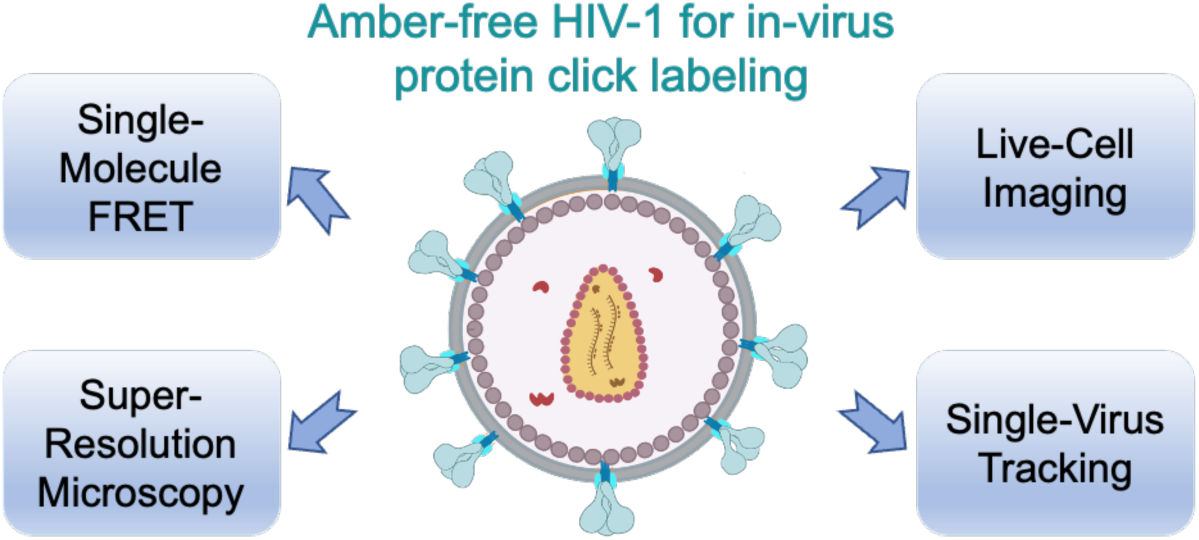

Amber-free HIV-1 infectious systems actualized minimal invasive Env tagging for smFRET, versatile for in-virus bioorthogonal click labeling in advanced microscopic studies of virus-host interactions.

## Introduction

Human immunodeficiency virus 1 (HIV-1) envelope glycoprotein (Env) is responsible for the host cellular receptor binding and primary neutralizing determinant. Thus, Env is one of the major targets for designing vaccines, antibodies, and antiviral drugs. Env interacts with the host cellular receptor CD4 and coreceptor CCR5/CXCR4 to mediate viral membrane fusion into target cells through a series of large-scale conformational changes^1–10^. Functional Env is a homotrimer-of-heterodimer consisting of gp120 exterior subunits and gp41 transmembrane subunits^11–16^. The gp120 subunit binds the primary receptor CD4 and coreceptor CCR5/CXCR4 on the susceptible target cells^4^. The binding of CD4 induces opening of gp120 and the formation of binding sites for coreceptor. After binding to CCR5/CXCR4 for gp120, additional conformational changes are relayed to the membrane-anchored gp41 subunit. Subsequent conformational refolding of gp41 induces fusion of the viral membrane with the host cell membrane^17–22^. Besides mediating viral entry, Env harbors virus-specific antigenic epitopes and thus serves as the main target of antibody responses during viral infection^1, 23–27^. Nevertheless, Env has evolved to successfully evade the host immune responses through multiple mechanisms, including extensive glycan shielding, regions of hypermutation, and conformational flexibility^28–31^. Collectively, Env exhibits a high capacity to undergo conformational switching events upon interacting with host cells, which are essential for virus entry and immune evasion during HIV-1 infection life cycle. Thus, a better understanding of conformational changes and dynamics of native Env on infectious viruses will inform the development of HIV-1 prevention and treatment strategies, such as vaccines, antibodies, and small-molecule inhibitors^1, 7–10^.

Single-molecule fluorescence resonance energy transfer (smFRET) imaging has provided unique insights into the HIV-1 entry mechanism by visualizing real-time conformational changes of Env in the context of intact virions, based on distance-sensitive energy transfer between a donor/acceptor pair of fluorophores labeled on Env (Figure 1A)^32–35^. Attaching donor/acceptor FRET-paired fluorophores to two sites of the host molecule within their Förster distance of a few nanometers is a prerequisite for applying smFRET^36^. We have previously used enzymatic labeling for smFRET imaging of HIV-1 Env^32, 37–39^, in which customized Cy3/Cy5 FRET-paired dyes are placed into Env gp120 variable loops V1 and V4, respectively, by enzymes that recognize introduced short peptides (Q3 and A1) and transfer dye-conjugated substrates to specific residues within these peptides^40, 41^. This approach results in the insertion of 6 and 12 amino acids into V1 and V4, respectively, of gp120. These insertions have generated concerns regarding the potential impact on structural integrity and the possible interaction with antibodies. Although these concerns have been addressed by extensive functional validations^32, 33, 42^, a minimal tagging approach for attaching dyes to Env is still desirable.

**Figure 1.**
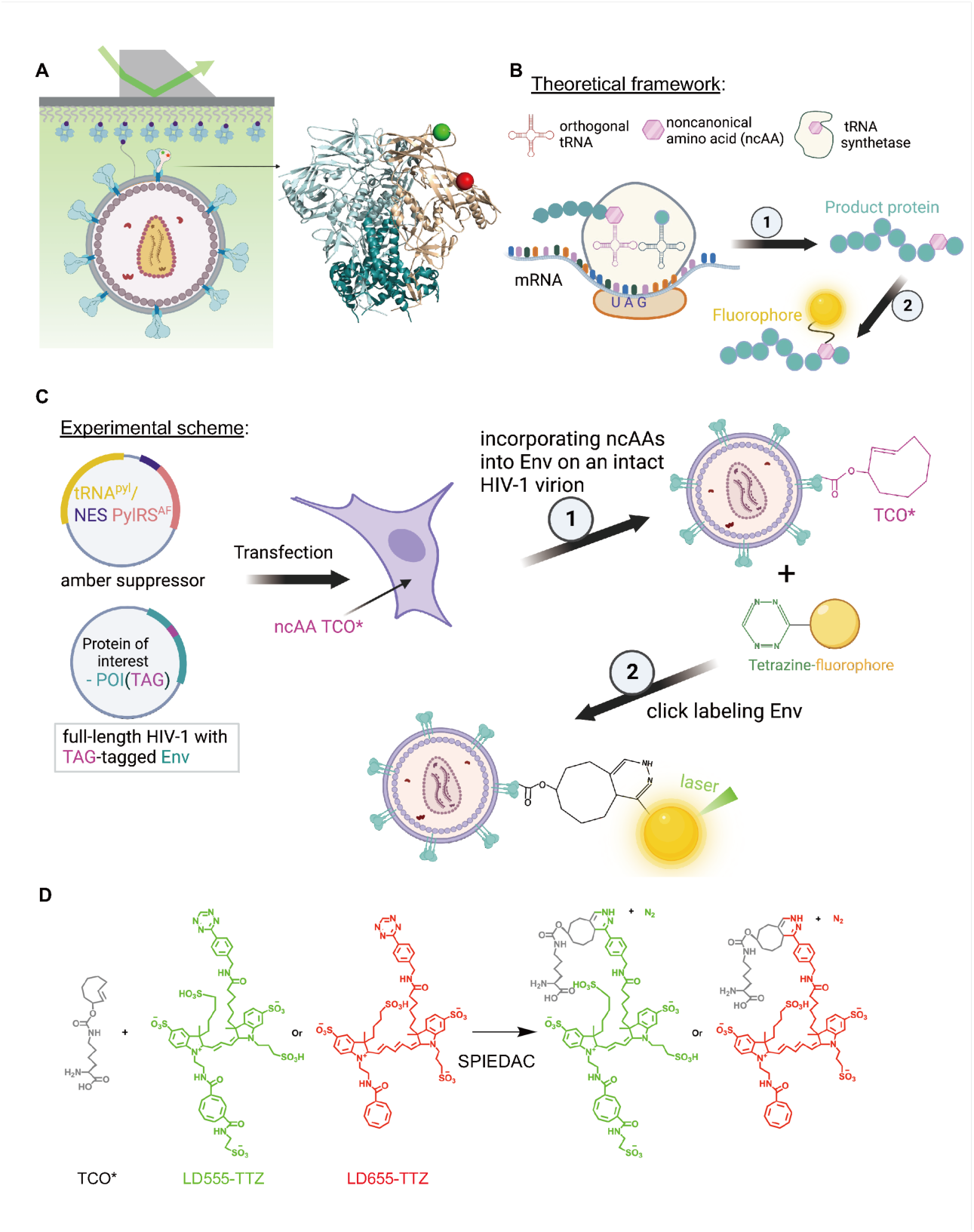
Experimental design for minimal invasive smFRET of HIV-1 Env on the intact virion using amber suppression and bioorthogonal click labeling. (A) smFRET of Env trimer in the virus context. An intact full-length HIV-1 virion carrying one Cy3/Cy5-paired labeled protomer within an Env trimer and elsewhere wildtype Env is immobilized on a quartz slide and imaged under the prism-TIRF microscope. The Env structure is adapted from PDB 4ZMJ (dually labeled gp120, brown; non-labeled gp120, cyan; gp41, blue). (B, C) Theoretical (B) and experimental (C) schematics of using two-step minimal genetically encoded amber (TAG) tags for fluorescence labeling - amber suppression (step **1**) followed by click chemistry (step **2**). Amber suppressor tRNA and tRNA synthetase (tRNA^Pyl^/NESPylRS^AF^) introduce ncAA trans-cyclooct-2-en – L – lysine (TCO*) into the desired TAG position in Env (POI) on an assembled intact virion generated in mammalian cells. The supply of TCO* to the transfected cells allows the incorporation of TCO* at the TAG insertion site in Env which can be clicked with fluorophores. (D) Bioorthogonal click labeling via SPIEDAC. Tetrazine-conjugated Cy3 and Cy5 derivatives (LD555-TTZ and LD655- TTZ) react with the strained alkene of the TCO* via strain-promoted inverse electron-demand Diels-Alder cycloaddition (SPIEDAC). Fluorophores/dyes are colored green and red, respectively.

Recently developed amber (TAG) stop-codon suppression through genetic code expansion in mammalian cells and copper-free bioorthogonal click chemistry to label dyes on proteins by only altering a single amino acid^43–49^ seems an ideal labeling strategy for Env. This approach of using the amber tag, advantageous over peptide tags, minimizes the impact of tagging on the structural and conformational integrity of proteins, thus offering a less invasive means for site-specific labeling Env (Table S1). Surprisingly, bioorthogonal click labeling of HIV-1 proteins in their close-to-native states has been rare, mainly with one report of labeling Env expressed on cells^43^ and the other of tracking capsids enclosed in viruses^50^. Both studies used laboratory-adapted virus/Env strain^43, 50^. Click labeling of HIV-1 proteins in the context of clinically relevant HIV-1 strains has been lacking. In the more relevant Env study of two reports mentioned earlier, a single dye coupled to an introduced amber in gp120 through amber-click two steps was achieved on cells with detectable Env expression, but no success was achieved with viruses due to incomplete suppression in mammalian cells and low copies of Env on each virion^43^. This approach includes translating ambers as clickable noncanonical/unnatural amino acids (ncAAs) by an engineered tRNA and its cognate tRNA synthetase (tRNA/RS) and the subsequent click labeling of dyes onto ncAAs^43, 51^. This strategy has been adapted to influenza A hemagglutinin^52^ and Ebolavirus glycoprotein^53^. However, coupling FRET-paired dyes to Env on viral particles through genetic code expansion–click chemistry has not been achieved, in part due to the increased structural complexity and lower copies of Env on HIV-1 compared to glycoproteins of influenza A or Ebolavirus in addition to the unfavorable incorporation efficiency of ncAA in mammalian cells.

Here we initiated the concept of amber-free HIV-1 to improve the amber-suppression efficiency of targeted Env proteins in mammalian cells and established an intact amber-free HIV-1 system for site-specific bioorthogonal click labeling by genetic code expansion. Using this system, we successfully incorporated dual ncAAs into Env on HIV-1 virions, which allowed click labeling for real-time smFRET imaging. smFRET results of click-labeled Env on the virus recapitulated the conformational landscapes of Env with different antibodies. The click-labeled Env has the potential to offer a more accurate and precise understanding of the conformational dynamics of Env in future studies. The amber-free HIV-1 and its derivative packaging systems provide new opportunities for fluorescence-based microscopic applications using minimally invasive bioorthogonal labeling in HIV-1 and other viruses.

## Results

### Construction of dually amber-suppressed click-labeled Env on intact HIV-1 virions

We aimed to design and establish a two-step minimally invasive labeling strategy for smFRET imaging on the virus under customized prism-based total internal reflection fluorescence (prism-TIRF) microscopy via minimal genetically encoding ncAA tags in gp120 and click labeling ncAAs with FRET-paired Cy3/Cy5 dyes (Figure 1). Incorporation of ncAAs into Env on intact HIV-1 virions can be achieved by co-transfecting HEK293T cells with plasmid tRNA^Pyl^/NESPylRS^AF^ and an amber-tagged Env plasmid, in the presence of ncAAs^43, 45, 54, 55^. We introduced ncAA trans-cyclooct-2-ene-L-lysine (The A isomer, from here on abbreviated TCO*) at an amber TAG stop codon engineered into gp120 by a pyrrolysine tRNA and its cognate aminoacyl tRNA synthetase (tRNA^Pyl^/NESPylRS^AF^) pair (Figures 1B, C). PylRS^AF^ is the double mutant (Y306A, Y384F) of the *M. mazei* pyrrolysine tRNA synthetase, accommodating bulky side-chain moieties in ncAAs. tRNA^Pyl^/NESPylRS^AF^ plasmid can be expressed in mammalian cells, such as HEK293T cells^44^. TCO* contains a trans-cyclooctene functional group, which reacts with tetrazine-conjugated Cy3 or Cy5 fluorophores by copper-free click chemistry via strain-promoted inverse electron-demand Diels-Alder cycloaddition (SPIEDAC, Figure 1D)^33, 43, 47^. tRNA^Pyl^/NESPyl RS^AF^ is an advanced amber suppressor system, which contains a robust nuclear export signal (NES) for improving amber-readthrough efficiency by relocating tRNA synthetase to the cytoplasm (where the translation occurs)^45^.

We next screened for suppression of Env TAG variants in the presence of tRNA^Pyl^/NESPylRS^AF^ in mammalian cells HEK293T by measuring the release of HIV-1_Q23 BG505_ (BG505 Env with Q23 virus backbone^56^) infectivity on TZM-bl cells. BG505 is a clinical Clade A transmitted/founder Env strain. We noticed that the HIV-1_Q23 BG505_ wildtype virus infectivity exhibited an approximate 50 to 100-fold reduction (Figures 2, S1) in the presence of the amber suppressor system (suppression condition) compared to the results under non-suppression condition (without the suppressor and TCO*). To identify amber-tolerable positions in gp120, we generated four full-length HIV-1_Q23 BG505_ Env TAG variants that each carried single TAG insertion at different sites within gp120 V1 loop, denoted as N133_TAG_, N136_TAG_, N137_TAG_, and T139_TAG_. The amber-inserted positions were the same or adjacent to the previously insertion sites of peptide tags^33, 42^. Among four variants, N136_TAG_ showed better amber tolerance (ncAA incorporation efficiency) than the other three, as indicated in relatively higher released infectivity under amber-suppression conditions (Figure S1). We combined N136_TAG_ with six amber insertion sites in the gp120 V4 loop and generated six dually amber-tagged HIV-1_Q23 BG505_ clones. Among six clones, N136_TAG_ S401_TAG_, and N136_TAG_ S413_TAG_ can better tolerate amber insertions in Env by showing vague signs of infectivity released under suppression conditions in contrast to under non-suppression conditions (Figure S1). Therefore, N136_TAG_ S401_TAG_ or N136_TAG_ S413_TAG_ Env variants likely can be used as candidates for achieving fluorescently dually click-labeled Env on intact viral particles for smFRET studies. We also noted the unsatisfying level of infectivity released from HIV-1_Q23 BG505_ wildtype and variants carrying TAG in Env, implying the need for improvements in amber suppression efficiency and its resulting ncAA incorporation to Env.

**Figure 2.**
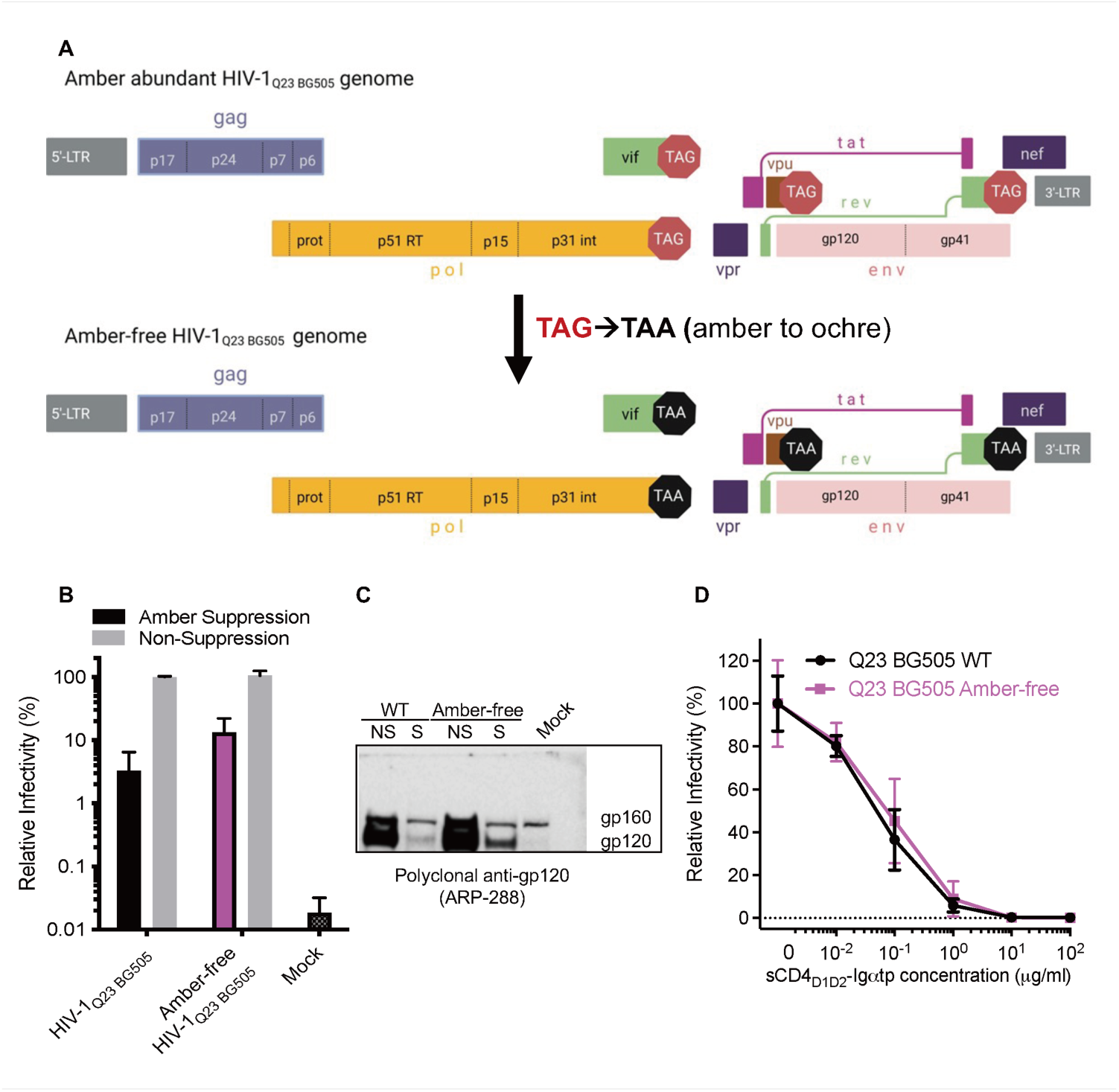
Construction and validation of an amber-free primary HIV-1 system. (A) Alternations of four amber TAG termination codons of pol, vif, vpu, rev proteins in full-length wildtype HIV-1_Q23 BG505_ to ochre TAA termination codons, generating an amber-free HIV-1 genome. (B–D) Amber-free HIV-1_Q23 BG505_ yields an approximate 10-fold increase in released infectivity (B) and an enhanced Env expression (C) and retains similar susceptibility to a very potent dodecameric sCD4_D1D2_ molecule (sCD4_D1D2_–Igαtp, D) compared to the wildtype. (B) Release of infectivity (mean ± SD) from HEK293T cells transfected with plasmids encoding amber-free or the wild-type HIV-1_Q23 BG505_ was measured on TZM-bl cells. Where indicated as “Amber-Suppression” or “S” a plasmid encoding tRNA^Pyl^/NESPylRS^AF^ was co-transfected with HIV-1 genome and the TCO* at 250 mM was added to the media. In contrast, only HIV-1 genome was transfected under the “Non-Suppression” or “NS” experimental condition (grey bar in B). WT or wildtype, the original amber-abundant HIV-1_Q23 BG505_. (C) Env processing and incorporation (Figure S1) for HIV-1_Q23 BG505_ were assessed by SDS-PAGE of virus supernatants, followed by western blotting using the polyclonal anti-gp120 and anti-HIV-IG. Repeated twice. (D) Neutralization curves (mean ± SD) are averaged from three independent experiments in duplicates.

### Development of an intact amber-free HIV-1 system for improving amber suppression efficiency in the target protein

To improve ncAA incorporation efficiency, we first explored why the released infectivity of HIV-1_Q23 BG505_ wildtype on target cells was 50 to 100-fold less under suppression conditions than under non-suppression conditions (Figure S1). By analysis of the complete genomic sequence, we observed the full-length HIV-1_Q23 BG505_ to be amber-abundant, containing four pre-existing amber termination codons in Pol, Vif, Vpu, and Rev open reading frames (Table S2), respectively. These amber codons might be translated to ncAA TCO* by amber suppressors against intrinsic cellular translation machinery. Consequently, the read-though of supposed-to-terminate-translation amber codons is problematic through the generation of additional amino acids at the C termini of the above proteins before termination (Table S2). Given the critical role of Pol in the HIV-1 infection lifecycle, our first attempt was made to change amber to oche (TAG to TAA) in Pol. Strikingly, a single TAG-to-TAA alternation in Pol improved the released infectivity under the suppression condition by approximately 5 to 10-fold (Figure S1), implying significant issues imposed by the pre-existing ambers in the HIV-1 genome. Thus, the previously observed unsatisfying efficiency of amber suppression and the resulting low ncAA incorporation into Env TAG variants were likely in part due to problematic error translations of pre-existing amber codons in HIV-1 and the competition of these ambers with artificially inserted ambers in the target protein Env.

To address the amber-abundant issues in HIV-1, we constructed an amber-free HIV-1_Q23 BG505_ system by modifying four termination codons from amber to ochre presented in the Pol, Vif, Vpu and Rev genes (Figure 2A). An approximately 10-fold increase in released infectivity was obtained in the amber-free HIV-1_Q23 BG505_ construct, compared with the amber-abundant one (Figure 2B). Env expression and incorporation into virions of amber-free and amber-abundant HIV-1_Q23 BG505_ constructs were further verified by immunoblots of virus supernatants using the polyclonal anti-gp120 and anti-HIV IgG (Human serum). Consistently, gp120 and gag P24 expression levels were higher in amber-free HIV-1_Q23 BG505_ than in the amber-abundant wildtype under the same suppression conditions (Figures 2C, S2). Amber-free HIV-1_Q23-BG505_ was highly sensitive to neutralization by a potent soluble dodecameric sCD4_D1D2_–Igαtp (12 × CD4), analogous to the sensitivity of wildtype one (Figure 2D). Thus, the developed amber-free HIV-1 system is an optimal backbone for inserting our genetically encoded amber tags in Env for smFRET studies.

### Intact amber-free HIV-1 carrying clickable dual ncAAs in Env retains neutralization sensitivities to trimer-specific neutralizing antibodies

We next evaluated the amber-suppressed efficiencies of previously screened Env TAG variants in our new amber-free HIV-1 system by measuring and normalizing the released infectivity. We identified thus far superior amber-tolerable positions: N136_TAG_ in the V1 loop, S401_TAG_ and S413_TAG_ in the V4 loop (Figure S1). Those positions were identical or adjacent to the previous Q3 tag or A1 tag insertion sites (Figure 3A). In combination with the Q3 tag in V1 (V1Q3) or the A1 tag in V4 (V4A1), we generated four dually tagged amber-free HIV-1_Q23 BG505_ constructs, including our primarily focused two dual-ncAA (or dual-amber) smFRET systems (N136_TAG_ S401_TAG_ and N136_TAG_ S413_TAG_) and two hybrid peptide/ncAA (or peptide/amber) systems (N136_TAG_ V4A1 and V1Q3 S401_TAG_) as comparisons (Figure 3A). Singly amber-tagged Env variants N136_TAG_, S401_TAG_, and S413_TAG_ retained ∼ 30% of infectivity activity (suppression efficiency) compared to the counterpart (Figure 3B, left), similar to the reported 5 ∼ 20% infectivity released from three different NL4-3 Env modified with single amber introduced into gp120^43^. Dually peptide/amber tagged N136_TAG_ V4A1 and V1Q3 S401_TAG_ showed ∼20% suppression efficiency (Figure 3B, middle). Dually amber-tagged Env (N136_TAG_ S401_TAG_ and N136_TAG_ S413_TAG_) were successfully suppressed with efficiencies of 10 to 20% relative to the wildtype counterpart (Figure 3B, right), comparable to the level of peptide/amber-tagged Env. Western blotting analysis of post-transfected cell supernatants revealed that N136_TAG_ S401_TAG_ and N136_TAG_ S413_TAG_ Env variants were successfully suppressed, proteolytically processed (gp120 detection), and incorporated onto amber-free HIV-1 virions, in contrast to no traceable Env expression under non-suppression conditions (Figure 3C, top). Both gp120 and gp160 expressions were observed in their post-transfected cell lysates (Figure 3C, bottom). The presence of gp160 in cell lysate and its absence in cell supernatant implies that precursor gp160 Env is largely (if not entirely) proteolytically cleaved as mature gp120/gp41 on the released HIV-1 virions from producer cells. Nanoparticle Tracking Analysis (NTA) showed the viral particles with a mode diameter size of 109.2 ± 15.9, 125.7 ± 3.4, and 121.8 ± 4.2 nm for respective amber-free wildtype, N136_TAG_ S401_TAG,_ and N136_TAG_ S413_TAG_ Envs-carrying intact HIV-1_Q23 BG505_ virus particles (Figure S1). The mode diameter was close to 120 nm in diameter size of natural HIV virions^57^, suggesting the normal assembly of ncAAs incorporated intact virions. We next tested the sensitivities of amber-free HIV-1_Q23 BG505_ N136_TAG_ S401_TAG_ and N136_TAG_ S413_TAG_ virus particles to trimer-specific neutralizing antibodies PG9, PG16, and PGT151, which target different epitopes on Env^33, 58, 59^. Neutralization cures of amber-free HIV-1_Q23 BG505_ N136_TAG_ S401_TAG_ and N136_TAG_ S413_TAG_ viruses resembled that of amber-free wildtype viruses (Figure 3D), indicating the preservation of neutralization sensitivities of dual-ncAA incorporated Env to trimer-specific antibodies. Thus, we demonstrated that our constructed dual-amber-tagged Env variants were successfully expressed with efficient ncAA incorporations, processed, and incorporated into intact amber-free HIV-1 viruses that retained neutralization sensitivities to Env-directed antibodies. These functional validations of viruses carrying two minimal genetically encoding tags in Env pave the way for minimally invasive smFRET imaging of Env.

**Figure 3.**
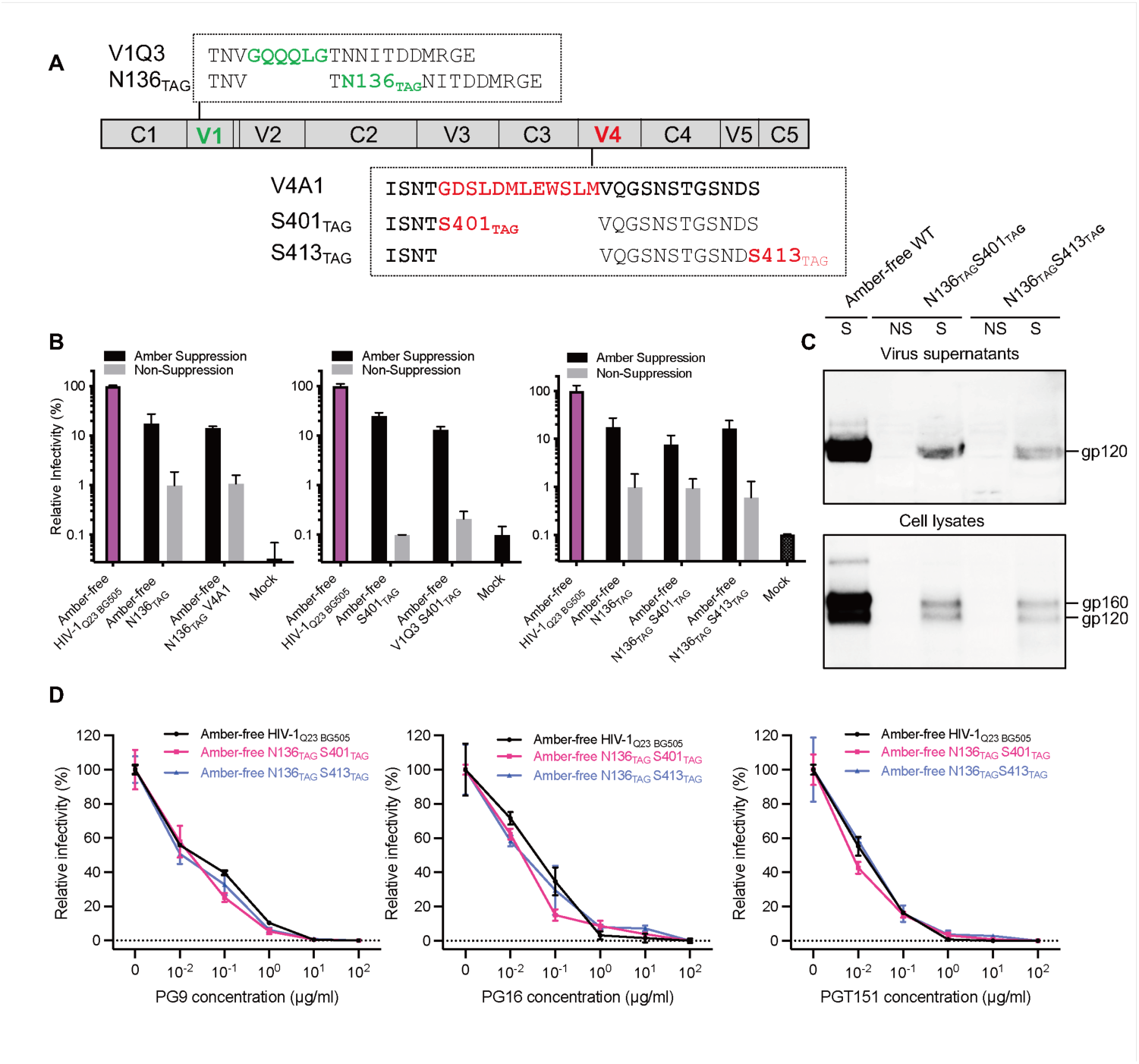
Efficient ncAA incorporation into dually amber-tagged Env on the amber-free HIV-1 virus that retains neutralization sensitivities to trimer-specific neutralizing antibodies. (A) Scheme showing tag insertion sites and denotations of labeling tags in BG505 gp120 subunit. Positions of optimized amber-tolerable sites and previously used peptide tags are in variable loops 1 and 4 (V1 and V4), respectively. Env with peptides (Q3 and A1) was denoted as V1Q3 and V4A1. Env tagged with ambers was denoted based on TAG insertion sites, such as N136_TAG_, S401_TAG_, and S413_TAG_. (B) Release of infectivity (mean ± SD) on TZM-bl cells from HEK293T cells transfected with plasmids encoding singly amber-tagged, dually peptide/amber-tagged, dually amber-tagged in Env_BG505_ in the context of an amber-free Q23 backbone (HIV-1_Q23 BG505_). (C) Env processing and incorporation of amber-free HIV-1_Q23 BG505_ and its dually amber-tagged Env (N136_TAG_ S401TAG and N136_TAG_ S413TAG) on the viruses from cell supernatants and lysates. Proteins were analyzed by immunoblotting using a polyclonal antiserum against HIV-1 gp120. (D) Neutralization of amber-free HIV-1_Q23 BG505_ and its dually amber-tagged Env by trimer-specific antibodies, PG9, PG16, and PGT151. mean ± SD. Dose-dependent relative infectivity was normalized to the mean value determined in the absence of the corresponding ligand.

### smFRET imaging of minimal genetically encoded amber-clicked Env in the amber-free HIV-1 context

We next established and performed smFRET imaging of fluorescent dual-ncAA-tagged Env (alternatively as amber-clicked Env) on amber-free intact virions (Figures 1A, 4). Where indicated, amber-clicked or dually amber-clicked Env simplifies the description of click-labeled Env generated by coupling Cy3/Cy5 derivatives to the TCO* molecules that are incorporated into Env via amber suppression. Similarly, the denotation of amber-click smFRET systems represents systems established based on amber suppression and click chemistry. Clickable dyes coupled to gp120 V1V4 loops, as exampled in N136_TAG_ S413_TAG_ (Figure 4A), include LD555-TTZ and LD655-TTZ. Two hybrid peptide/ncAA FRET systems are used as comparisons, in which Envs are enzymatically labeled in combination with click labeled with paired dyes. Enzymatically labeled dyes include cadaverine-conjugated Cy3 and coenzyme A (CoA)-conjugated Cy5 derivatives, similar to previously used dyes^32, 33, 42, 60^. For smFRET imaging, intact HIV-1 virions were produced by co-transfecting HEK293T cells with separate plasmids encoding the amber-free HIV-1_Q23 BG505_ carrying wildtype Env or tagged Env variants, and the suppressor tRNA^Pyl^/NESPylRS^AF^, supplemented with the TCO* added to the media at 250 µM. As smFRET monitors a single FRET-labelled gp120 protomer within an Env trimer in otherwise wildtype trimers on a single virion, we could achieve 1labeled-protomer:1trimer:1virion by adjusting the ratio of tagged vs. wildtype Env^32, 33, 61^. Based on the double-amber suppression efficiency, we use an approximate 4:1 ratio of amber-free HIV-1_Q23 BG505_ wildtype and HIV-1_Q23 BG505_ N136_TAG_ S401_TAG_ or N136_TAG_ S413_TAG_ to achieve ideal conditions for smFRET: single FRET-labelled gp120 protomer embedded in an otherwise wildtype environment. Analogously to the previously described^33^, LD555-TTZ and LD655-TTZ could be coupled randomly to the two ncAAs (TCO*) in Env presented on the intact amber-free intact HIV-1 virion. The virions were further immobilized on a streptavidin-coated quartz plate-coverslip sample chamber through the interaction between the virus membrane and biotinylated lipids. LD555-TTZ labeled on the immobilized virions were excited by a 532-nm continuous wave laser, The fluorescence intensities of donor and acceptor dyes (LD555-TTZ and LD655-TTZ) were recorded simultaneously for a period of 80 seconds at 25 Hz by our customized prism-TIRF microscopy. We extracted hundreds of fluorescence traces with anti-correlated features between donor and acceptor fluorescence intensities (Figures 4B, C; top). Anti-correlation is the revealing signature of energy transfer between donor and acceptor, reflecting conformational changes of Env trimers on individual virions. The donor or acceptor photobleaching point, either acceptor bleaching first (Figure 4B) or donor bleaching first (Figure 4C), indicates the detection of a single FRET-labeled Env on an immobilized intact virion. We further derived the FRET efficiency (FRET traces/trajectories, exampled in Figures 4B, C; bottom), documenting real-time conformational changes over time of individual Env on virions.

**Figure 4.**
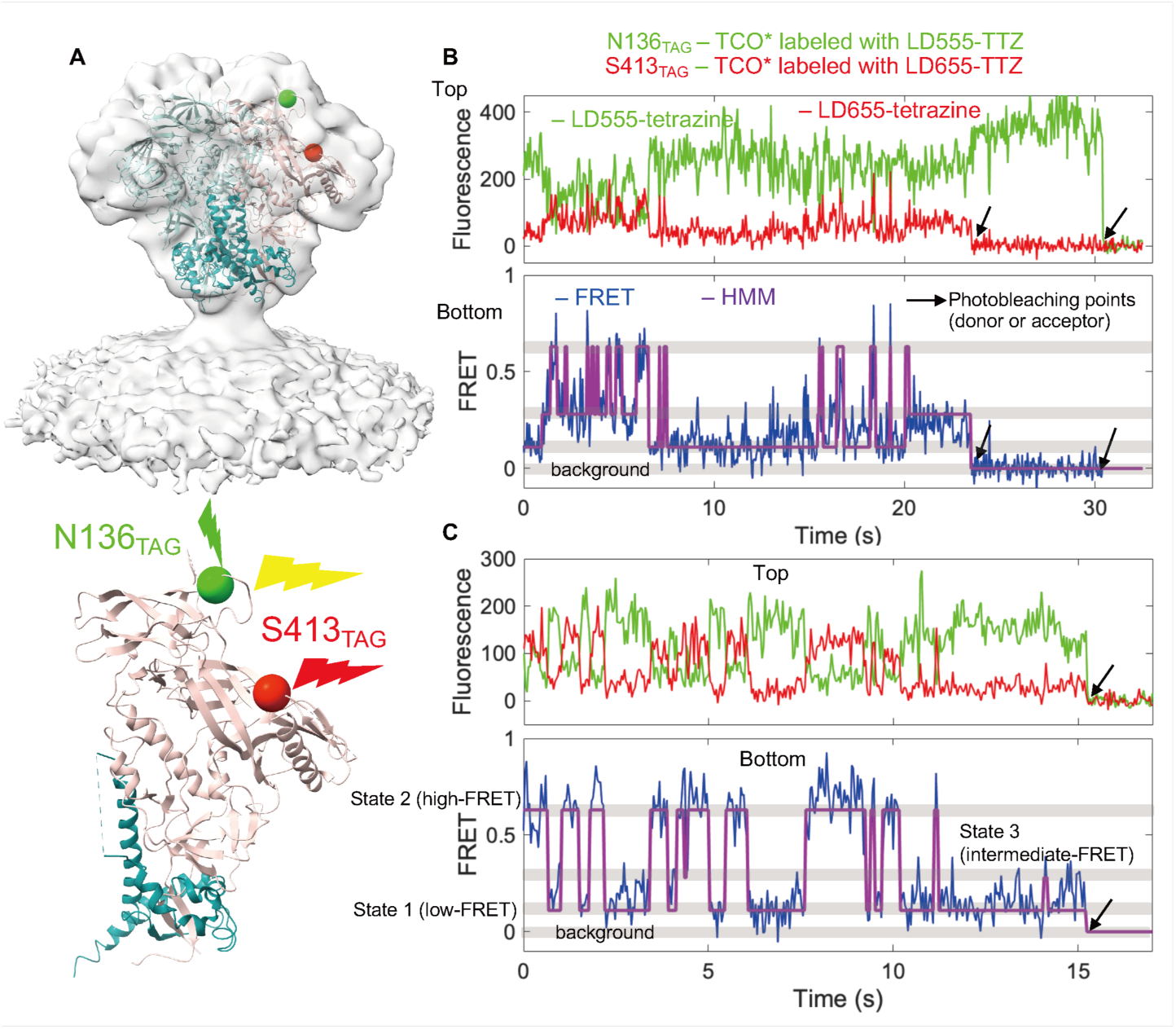
Representative fluorescence and FRET traces for unliganded dually amber-click labeled Env on intact virions at the single-molecule level. (A) A donor/acceptor-paired labeled protomer within an Env trimer at indicated sites on an intact virion was immobilized on a quartz slide and imaged by prism-TIRF microscope. Donor dye was excited with an incident laser and transferred its energy to nearby acceptor dye depending on its proximity. The structure of the Env trimer (PDB 4ZMJ)15 was fitted into *in-situ* electron density map of the ligand-free trimer (EMD- 21412)34. Dually labeled gp120 monomer, brown; unlabeled gp120 monomer, cyan; gp41, blue; donor fluorophore, green; acceptor fluorophore, red. (B, C) Two example fluorescence traces (Top panel) and derived FRET traces (Bottom panel) of individual dually amber-suppressed click-labeled Env_BG505_ on intact amber-free HIV-1_Q23_ virions. LD555-TTZ and LD655-TTZ were attached to TCO* at sites located in N136_TAG_ and S413_TAG_, respectively (Top panel). Their corresponding FRET efficiency values (FRET) trajectory (blue) with overlaid idealization were generated using Hidden Markov Modeling (HMM) (purple) (Bottom panel). The monitored individual gp120 monomers undergo transitions between three FRET-identified conformational states (State 1, low FRET; State 2, high-FRET; State 3, intermediate-FRET). Photobleaching points (B, the acceptor bleaches first; C, the donor bleaches first) indicate time-course fluorescence/FRET monitoring at the single-molecule level.

### Recapitulation of Env conformational profiles by minimally invasive amber-click smFRET systems in the context of intact amber-free virions

Qualitative and quantitative smFRET analysis of amber-clicked Env in the amber-free virus context revealed globally consistent three-state conformational profiles of Env on the viruses with results obtained from previous dual-peptide-tagged and two hybrid peptide/ncAA-tagged Envs (Figures 5, S5, Table S3). Our previous smFRET studies using peptide tags have shown that native Env on the virus is intrinsically dynamic, transiting from the pre-triggered conformation (State 1) through a partially open intermediate (State 2) to the fully activated open conformation (State 3) upon activation by CD4 molecules^32, 33, 38, 42^. FRET traces/trajectories report time-correlated relative donor-acceptor distance changes associated with conformational changes between V1 and V4 regions within the Env monomer. Compiling hundreds of FRET traces obtained under different conditions led to respective merged histograms fitted with Gaussian functions, indicating conformational distributions with state occupancies (Figures 4, 5). FRET histograms of Env revealed by dually amber-clicked HIV-1_Q23 BG505_ N136_TAG_ S401_TAG_ (Figures 5A-C), N136_TAG_ S413_TAG_ (Figure 5D-F), as well as mix-labeled N136_TAG_ V4A1 or V1Q3 S401_TAG_ (Figure S1), were in good agreement with previous results from dual-peptide V1Q3 V4A1 FRET system (overlaid lines). Similarly, the three-state conformational ensemble contained State 1 (low-FRET with mean 0.11 ± 0.02), State 2 (high-FRET with mean 0.63 ± 0.05), and State 3 (intermediate-FRET with mean 0.28 ± 0.04). Transitions between low- and high-FRET states and high- and intermediate-FRET states occurred frequently, but the low- and intermediate-FRET states were rarely observed, as shown in the representative traces (Figures 4B, C, and S5A). The unliganded N136_TAG_ S401_TAG_, N136_TAG_ S413_TAG_, N136_TAG_ V4A1, and V1Q3 S401_TAG_ tagged Env in the amber-free virus context all predominantly resided in a low-FRET State 1 conformation with the mean occupations of 41%, 50%, 42%, and 52%, respectively, similar to the previous studies^33, 42^ (Figures 5, S5, Table S3). Consistent with the previous observations, potent sCD4_D1D2_–Igαtp shifted the conformational landscape of Env towards intermediate-FRET State 3-dominant distributions with the occupations of respective 38%, 47%, 45%, and 50% from the above-mentioned four smFRET systems (Figures 5, S5, Table S3). Similarly, adding the PGT151 antibody that asymmetrically targets the interface between gp120 and gp41 resulted in the preferential conformation to be high-FRET State 2 with the mean occupations of 54%, 58%, 57%, and 50% (Figures 5, S5, Table S3). Donor/acceptor dyes coupled to distinct pairs of sites on Env may contribute to the observed variations in the relative occupancy of three conformations in different FRET systems. More importantly, the trending of dominance across all established FRET systems remained the same under different ligand-stabilizing conditions (Figure 5G). This consistency reinforced our previous finding of Env sampling three primary large-scale global conformations. Altogether, minimally invasive amber-click smFRET systems of Env in the context of intact amber-free virions recapitulated the previously delineated conformational landscape of Env and the corresponding responses to conformation-stabilizing ligands.

**Figure 5.**
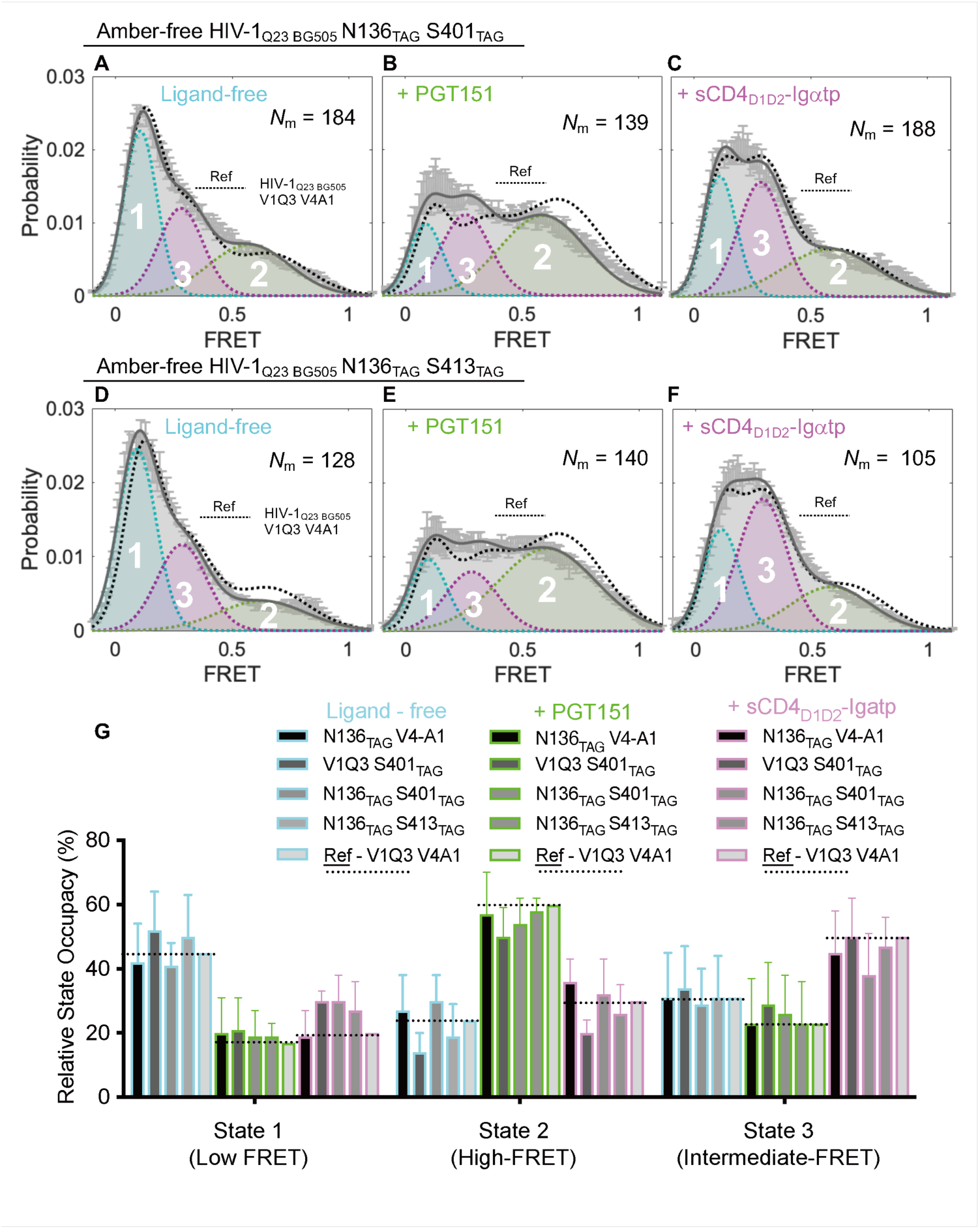
Recapitulation of three-state conformational landscapes of Env by dually amber-suppressed click-labeled smFRET systems on the virus. (A–C) FRET histograms of dually click-labeled HIV-1_Q23 BG505_ at sites N136_TAG_ S401_TAG_ under different conditions: ligand-free (A), PGT151-bound (B), and sCD4_D1D2_–Igαtp-bound (C). The number (*N*_m_) of individual dynamic traces was compiled into a conformation-population FRET histogram and fitted into a 3-state Gaussian distribution (solid gray line) centered at ∼0.1-FRET (dashed cyan), ∼0.3-FRET (dashed red), ∼0.6-FRET (dashed green). The fitted distributions (solid gray line) were overlaid with the corresponding referenced distributions of enzymatic-labeled HIV-_1Q23 BG505_ V1Q3 V4A1 (dashed black lines), where ref stands for the referenced distribution. (D–F) Experiments as in A–C, revealing conformational landscapes for the ligand-free (D) and upon binding of PGT151 (E) or sCD4_D1D2_–Igαtp (F) to dually amber-suppressed click-labeled HIV-1_Q23 BG505_ N136_TAG_ S413_TAG_. (G) Bar graph of relative conformation-occupancy of virus Env using different FRET probes. FRET histograms represent mean ± SEM, determined from three randomly assigned populations of FRET traces under indicated experimental conditions. Relative occupancies of each conformational state and fitting parameters were summarized in Table S3.

## Discussion

Here, we developed and optimized an intact amber-free HIV-1 system that significantly improved the incorporation of minimally invasive tags (ncAAs) at introduced ambers into Env at single-residue precision by genetic code expansion. The dually amber-tagged Env trimers were efficiently suppressed and successfully incorporated into viral particles that largely retained viral infectivity and completely resembled neutralization sensitivities to three tested trimer-specific neutralizing antibodies. These advancements further actualized bio-orthogonal click labeling of Env with click-chemistry-reactive dyes and the subsequent establishment of the minimal invasive amber-click smFRET systems of Env on gp120 variable loops. The amber-free HIV-1 system is versatile and valuable for in-virus protein biorthogonal click labeling.

### Demands for minimally invasive labeling strategy in smFRET imaging of Env

Site-specifically fluorescence labeling of HIV-1 Env close to its native state without compromising its functionality is highly desired but challenging. Variable loops of gp120 were shown to be the most tolerant sites for fluorescent protein insertions^62^. Insertions of optimized EGFP into the variable loops of Env can partially retain Env functions^62, 63^, but they are unsuitable for studying the conformational dynamics of Env. The conventional way to label proteins for smFRET is to introduce cysteines based on maleimide-thiol chemistry^64^. A considerable number (several dozen) of preexisting essential cysteines on Env trimer make the maleimide-thiol approach infeasible. To achieve smFRET of Env, we had previously deployed an alternative approach of introducing short peptide tags (Q3 and A1)^40, 41^ into variable loops of gp120^32, 33, 42^. smFRET imaging of gp120 within Env trimers on the surface of intact HIV-1 virions has granted exclusive access to dynamic aspects, conformation-associated immune-evasive profiles, antagonism by fusion inhibitors, and, importantly, spike-based vaccine design of HIV-1 Env^32–34, 38, 42, 65–67^. Nevertheless, the introduction of a few amino acids still holds the possibility of negative impacts on accurately revealing Env conformation in unpredictable ways. For instance, reasonable concerns have been raised regarding the potential interactions between antibodies and introduced peptide tags. Amber-click labeling (Table S1) offers possibilities of site-specific, minimally invasive, fluorescent-labeling for Env at single-residue precision by only altering one single residue for each tag insertion^51, 68^. Thus, it is the method of choice for staining Env within its variable loops of gp120 and even promising for accessing the sterically tight gp41. The strategy for incorporation of ncAAs facilitates identifying more permissive labeling sites that lower the risk of functional impact^69^. Therefore, labeling Env using amber-click strategy would be highly advantageous for the Env studies using smFRET and other advanced fluorescence microscopy techniques.

### Challenges in amber-click labeling HIV-1 proteins on the virus, especially for the naturally sparse Env

To the best of our knowledge, there have been no reports of site-specifically amber-suppressed click labeling (amber-click) of Env or dually click labeling of HIV-1 proteins on intact HIV-1 virions. Given the rapidly increasing number of applications of amber-click fluorescence labeling of proteins in living cells, neurons, and viruses^43, 46, 55, 70–72^, this is somewhat surprising. It may, however, be somewhat expected considering one of the major limitations in the genetic code expansion approach, intrinsically incomplete suppression in mammalian cells^43, 54, 73^. Two pioneering attempts have been made in lab-adapted HIV-1, including one of visualizing Env on cells ^43^ and the other of tracking capsid on viruses^50^. Sakin and colleagues have generated single amber-click labeled Env expressed on mammalian cells for living-cell and super-resolution imaging of Env mobility at the plasma membrane^43^. However, this pioneered study did not achieve click labeling Env on the viral particles due to low single ncAA incorporation into Env at each of the three optimized sites^43^. Their results are completely in line with our previously inefficient amber suppression of Env (Figure S1) and a lack of apparent Env incorporation on viral particles in immunoblots (unpublished). Singly click labeling of capsid has been achieved until recently, in which live-cell imaging and super-resolution microscopy of click-labeled capsid enabled visualizing/tracking capsid on lab-adapted HIV-1_NL4-3_ viruses during the route from the plasma membrane to the nuclear^50^. The use of primary HIV-1 isolates rather than lab-adapted ones could be more informative. Of note, the dramatic difference in the number of capsids (thousands of copies) versus Envs (a few copies)^13, 74–76^ in/on each virion may explain the success of amber-click labeling capsid on the virus^50^ but not yet for Env^43^. As singly amber-click labeling of HIV-1 proteins in the virus context has proven to be challenging, the hurdles to success in dually click labeling of Env are expected to be exponential. Major challenges of low copies of Env on HIV-1, incomplete amber suppression in mammalian cells, and high Env structural complexity have been predictably impeding our long-sought-after smFRET imaging of Env on the virus.

### Pinpointing a limiting factor of amber suppression in the wildtype HIV-1 genome

We found an intrinsic issue of abundant amber termination codons in the HIV-1 genome. We hypothesized that preexisting ambers in the HIV-1 genome could result in error translations in viral proteins by incorporating ncAAs into amber codons against the internal cellular translation machinery. To test our hypothesis, we constructed an intact amber-free HIV-1 system by genetically modifying amber stop codons (TAG) to ochre stop codons (TAA) in critical protein-coding gene sequences, warranting their correct translating terminations. By replacing four nature ambers in HIV-1 primary isolate Q23 to ochre, we rescued virus infectivity by more than 5 to 10-fold from the artificially introduced amber translational machinery (Figures 2A, B). Consistent results were also seen in a substantial improvement of Env incorporation into viral particles and the retainment of neutralization sensitivity to potent 12xCD4 molecules (Figures 2C, D). Changing the amber stop codon of Vpu to ochre causes a point mutation in Env, but it is within the signal peptide and does not appear to interfere with Env secretion (Figure 2C). Modifying the amber of Rev to ochre results in a mutation in the C-tail of Env gp41, but it does not seem to affect Env conformational dynamics (Figure 5, S5). The abnormal extension of Pol is likely the most serious problem for amber suppression of naturally amber-abundant HIV-1_Q23_ (Table S2). We demonstrated that the previously observed 50 to 100-fold reduction in virus infectivity in the presence of amber suppressors was likely attributed to two major factors: the error translation of naturally existing amber termination codons in both HIV-1 genome (especially the abnormal translation of Pol) and mammalian cells. Despite the unsolved incomplete suppression issue in mammalian cells, the constructed amber-free HIV-1 provides an optimized system for screening tolerable amber labeling sites within Env and other HIV-1 proteins on viral particles.

### The amber-free HIV-1 system enables dually amber-suppressed click-labeled Env on the virus, enabling minimal invasive *in situ* smFRET imaging of Env

We further contextured amber-tagged Env in our newly developed amber-free HIV-1 system and successfully optimized two candidate clones encoding dually TAG-tagged Env at two pairs of sites. In contrast to the pioneered study^43^ and our previous attempts, our infectivity and immunoblot results provided more apparent evidence of enhanced amber suppression efficiency and resulting effective ncAA incorporation in Env (Figure 3). Produced viruses carrying dual-ncAAs incorporated Env retained neutralization sensitivities to three trimer-specific neutralizing antibodies, including Env apex-targeting PG9 and PG16 and gp120/gp41 interface-directed PGT151 (Figure 3). The consistent immunoblots and neutralization results indicate efficient trimerization and maturation of amber-tagged/ncAA-tagged full-length Env proteins, which present properly on intact HIV-1 virions. Results from the validation checkpoints, as mentioned earlier, imply that our two clickable ncAA-tagged Env variants for smFRET imaging largely resemble wildtype Env with minimal effect on functionalities. smFRET results of dually amber-clicked Env in parallel with hybrid peptide/amber Env were in global agreement with our previous observations^32, 33, 42^ (Figures 4, 5). The three-state conformational landscape of ligand-free Env was captured, and so did the conformational responses to open-trimer-stabilizing sCD4_D1D2_–Igαtp and asymmetric-trimer-stabilizing PGT151. These results validated our previous findings of three-state conformational distributions and shifts revealed by smFRET of peptide-tagged Env^32, 33, 42^, further ruling out the scenarios that sCD4_D1D2_–Igαtp and PGT151 affect peptide tags/dyes. The amber-click smFRET systems of Env gp120 is an advanced version of our extensively used enzymatic system. The amber-free context also holds great promise for smFRET imaging of the highly constrained Env transmembrane subunit gp41 as fusion machinery, of which conformational events remain elusive. The sterically tight gp41 cannot tolerate 6-8 amino acid peptide tags deployed for smFRET imaging of gp120, despite numerous attempts made by us and our smFRET peers. The amber-free HIV-1 system is expected to significantly increase our success rate of site-specifically introducing a pair of clickable donor/acceptor dyes into conformational switching regions of gp41 by only altering a single amino acid per site. Therefore, the potential of amber-click Env in the amber-free context will be exploited and implemented in our following smFRET imaging of conformational dynamics and allostery during virus entry.

### Application of an amber-free HIV-1 infectious system for in-virus protein bioorthogonal labeling

The amber-free HIV-1 infectious system developed in this study allows for broad in-virus protein bioorthogonal labeling, which is compatible with advanced fluorescence microscopic/nanoscopic imaging (Figure 6). The amber-free clinically relevant infectious virus system is highly versatile for screening amber-tolerable sites in any HIV-1 proteins, offering a high chance or efficiency of labeling proteins in/on HIV-1 virions. It, therefore, permits visualizing/tracking under-investigated bioorthogonally labeled viral proteins within the context of infectious virus particles close to their native states during the trafficking of viruses to live cells using live-cell microscopy, correlative imaging, and super-resolution microscopy (Figure 6). For example, the identified amber-tolerable N136_TAG_, S401_TAG_, and S413_TAG_ sites on Env in the amber-free Q23 virus context can be used for virus tracking on living cells using confocal, TIRF, super-resolution microscopy, or single-virus tracking. Searching for bioorthogonal clickable sites in the capsid in the context of our amber-free clinically relevant HIV-1_Q23_ infectious system, a step further from using lab-adapted HIV-1 isolate^50^, is promising for tracking capsid uncoating and other imaging applications. To better reap the benefits from superior amber suppression efficiently in our amber-free virus context, we also made other versions of amber-free HIV-1 systems. These systems include the replication-incompetent version with RT (reverse transcriptase) impaired (ΔRT), a new virus packing system with BG505 Env deletion (ΔEnv), and a double impairment/deletion of RT and Env (ΔRT ΔEnv). The ΔEnv packaging systems are applicable to make pseudo-typed viruses carrying other Env strains or surface glycoproteins of other enveloped viruses (analogously to Env of HIV). We will share these clones with our peer scientists to make the best use of viral protein bioorthogonal labeling in the context of our amber-free replication-competent or replication-incompetent HIV-1 systems.

**Figure 6.**
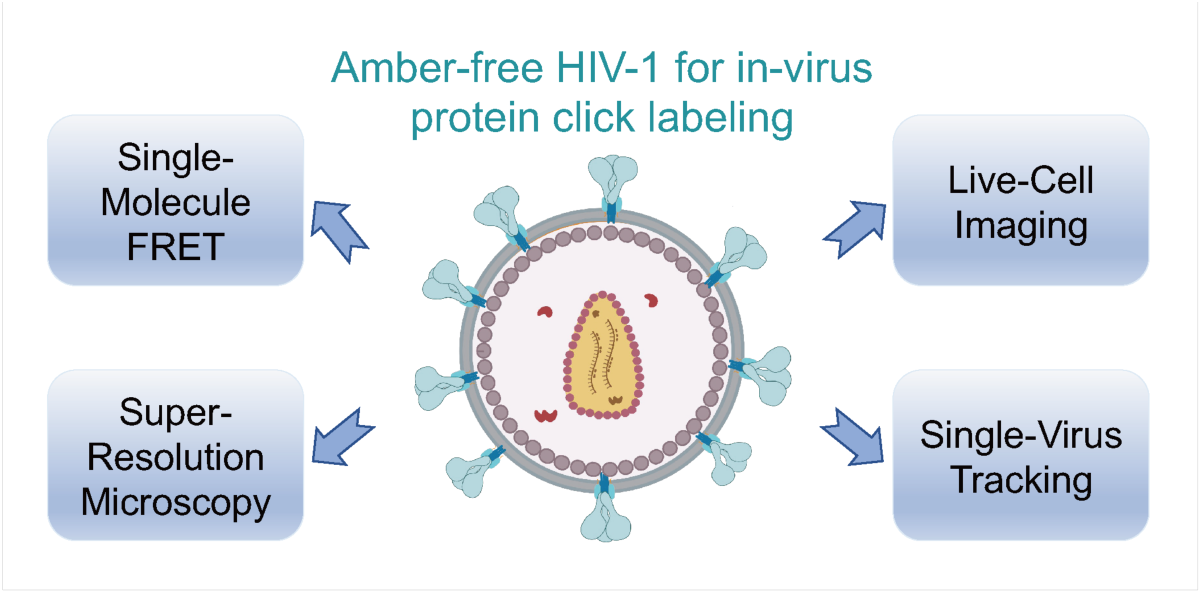
The broad microscopic applications of amber-free clinically relevant HIV-1 infectious systems for in-virus protein bioorthogonal click labeling in studying virus entry, trafficking, and egress in/on living cells. The amber-free replication-competent/incompetent and Env-truncated HIV-1 clones developed in this study are highly versatile for screening amber-tolerable sites in any HIV-1 proteins and even Env-analogous surface glycoproteins of other viruses.

### Limitations of this study

The amber-suppressed efficiencies of Env are still not perfectly satisfactory, even with the advanced amber-free HIV-1 system. Two factors likely dominate this issue. Firstly, the efficiency of incorporating ncAAs is still limited by unpredictable competition with cellular amber codons translations e.g., release factors^43, 72, 77–80^. Secondly, the tRNA^pyl/^PylRS^AF^ and its advanced NES systems from Methanosarcina species are not exploited fully in the eukaryotic cells^43^, leading to the decrease of amber-suppression efficiency with the expectation of the exponential decrease for dually amber-suppression in the same protein^81^. Future efforts to develop promiscuously amber-suppressed systems with higher suppression efficiency are thus required, e.g., improved pylRS system with an enhancement of translations^45^, and designed orthogonal tRNA/pylRS pair permitting simultaneous incorporation of two distinct unnatural amino acids against different stop codons^44^. Due to the instrumental limitations, we have not yet explored their full potential in advanced microscopic studies of click-labeled viral proteins beyond smFRET imaging of Env. Nevertheless, the success of dually click labeling Env at the single-molecule level in this study holds great promise for bioorthogonal click labeling other viral proteins with high copies at the ensemble or single-molecule levels.

In conclusion, we achieved dually site-specific incorporation of clickable unnatural amino acids into Env on intact amber-free HIV-1 virions by genetic code expansion. The amber-free system could open new opportunities for biorthogonal click labeling of functional proteins in HIV-1 and Env-analogous spike glycoproteins of other viruses.

## Author contributions

M.L., Y.A., and J.R.G. conceived this study. M.L., J.R.G., and Y.A. performed mutagenesis. Y.A. and M.L. designed and performed virus infectivity/neutralization assays. Y.A., Y.H., G.Z., and W.Q. performed immunoblotting experiments. Y.H., G.Z., D.G., and A.H. prepared plasmids and cultured cells. M.L., Y.A., and Y.H. prepared and purified fluorescently labeled viruses, and performed smFRET experiments and data analysis. Y.A., Y.H., and M.L. made figures, tables, and schematics. E.A.L and R.B provided analytical tools and reagents. B.Z., J.A. and P.D.K provided antibodies and other ligands, including the potent dodecameric soluble CD4 molecules. Y.A. and M.L. wrote the manuscript with inputs from P.D.K., E.A.L., and J.R.G. Correspondence and requests for materials and data should be addressed to M.L.

## Acknowledgments

## Acknowledgements

We thank the HIV Reagent Program (Division of AIDS, NIAID, NIH) for the neutralizing antibodies PG9 and PG16. PG9 and PG16 were obtained through the NIH HIV Reagent Program, Division of AIDS, NIAID, NIH: Anti-Human Immunodeficiency Virus 1 gp120 Monoclonal Antibody PG9 (ARP-11557) and PG16 (ARP-12150), contributed by International AIDS Vaccine Initiative. We thank Dr. Walther Mothes for his advice and the support from his laboratory, where some of these experiments were initiated. We thank Dr. L. Vijaya Mohan Rao and Dr. Torry Tuck for all their continued support and advice. We are grateful to Dr. Scott Blanchard for sharing smFRET software and the design of the prism-TIRF microscope. We thank Dr. David Derse for sharing HIV In-GLuc plasmids. We thank Polyhedron Optics LLC for the consulting service regarding the customized prism-TIRF microscope.

## Data availability

The data that support the findings of this study are available from the corresponding author upon reasonable request.

## Supporting Information

This file includes:

Materials and Methods
Figures S1 to S5
Tables S1 to S3

### Materials and Methods

#### Cell lines

HEK293T cells (human embryonic kidney, ATCC) were used for producing replication-complement and replication-incompetent HIV-1 viruses. TZM-bl cells (generated from HeLa cells) that stably express receptor CD4 and co-receptors CCR5 and CXCR4 were used as target cells for quantifying the infection abilities of produced HIV-1 viruses^82^. These cell lines were cultured in high glucose Dulbecco’s Modified Eagle Medium (DMEM, Gibco) supplemented with 10% (v/v) fetal bovine serum (FBS, Gibco), 100 µg/mL penicillin-streptomycin (PS, Gibco), and 6 mM L-glutamine (Gibco), and maintained in a 37 °C incubator supplied with 5% CO2.

#### Plasmid construction

The wildtype HIV-1_Q23 BG505_, an infectious clade A clone, contains full-length Env_BG505_ with Q23 as the backbone. HIV-1_Q23 BG505_ ΔRT is a replication-incompetent clone that lacks the reverse transcriptase gene^32, 33, 42^. Amber-free HIV-1_Q23 BG505_ plasmid was generated from HIV-1_Q23 BG505_ plasmid in which TAG stop codons presented in the Pol, Vif, Vpu and Rev genes were altered to TAA stop codons. Tagged-Env_BG505_ plasmids with amber-free Q23 as backbone include amber-free HIV-1_Q23 BG505_ N136_TAG_V4A1, V1Q3S401_TAG_, N136_TAG_S401_TAG_ and N136_TAG_S413_TAG_. Amber-free N136_TAG_V4A1 plasmid was tagged with a single amber codon TAG at the position of codon 136 (Asn) in the variable V1 loop and an A1 peptide tag (A1, GDSLDMLEWSLM) in the variable V4 loop within gp120 of Env, while plasmid amber-free V1Q3 S401_TAG_ contained the insertions of Q3 peptide tag (Q3, GQQQLG) in V1 and a single amber codon at the position of 401 (Ser) in the V4 of gp120. Similarly, N136_TAG_S401_TAG_ contains two ambers inserted at positions 136 (Asn) in V1 and 401 (Ser) in V4, respectively. N136_TAG_ S413_TAG_ was tagged by two ambers introduced at positions 136 (Asn) in V1 and 413 (Ser) in V4, respectively. These plasmids were constructed from amber-free HIV-1_Q23 BG505_ and HIV-1_Q23 BG505_ ΔRT for generating amber-click labeled Env in the context of intact virions. During the initial screening of amber-tolerance sites on Env_BG505_ V1 and V4 loops, single amber- or dual amber-tagged Env were generated from wildtype amber-abundant HIV-1_Q23 BG505_. We generated different versions of amber-free HIV-1_Q23 BG505_ clones, including ΔRT, ΔEnv (a new virus packing system with BG505 Env deletion), and a ΔRTΔEnv. All the plasmids were sequenced before being used.

#### Viral infectivity measurements

Viral infectivity of HIV-1_Q23 BG505_ was determined by using a Gaussia luciferase reporter^83, 84^ and quantified using a Gaussia luciferase flash assay kit (ThermoFisher Scientific, cat#16158), according to the manufacturer’s instructions. Briefly, amber-abundant or amber-free HIV-1_Q23 BG505_ viruses, or Env-tagged ones were produced, respectively, in HEK293T cells that were co-transfected with the corresponding HIV-1_Q23 BG505_ plasmid, and intron-regulated luciferase plasmid (HIV-1-InGluc) using transfection reagent polyethyleneimine (PEI Max^TM^, polysciences) or Fugene 6 (Promega, cat#E2311). Under amber suppression conditions, in addition to the HIV-1_Q23 BG505_ and HIV-1-InGluc plasmids, the transfection solution also includes a tRNA^Pyl^/NESPylRS^AF^ plasmid (1:3 HIV- 1_Q23 BG505_ plasmid) supplemented with 250 µM unnatural amino acid (ncAA) transcyclooct-2-ene lysine (TCO*, Sichem). Viruses in the cell supernatants were harvested at 40 hour (h) post-transfection, filtered through 0.45 μm filters (Pall Corporation), and concentrated by ultracentrifugation at 25,000 rpm for 2 h. Viruses diluted in the medium were added onto TZM-bl cells pre-seeded in a 48-well plate. At 40 to 48 h post-infection, 100 µl cell supernatant was transferred to 96-well plates for the measurement of gaussian luciferase activity using a Gaussia luciferase flash assay kit. Relative viral infectivity was determined by normalizing relative light units (RLU) to the control group in each assay.

#### Immunoblot assays

Viral particles of amber-abundant, amber-free HIV-1_Q23 BG505_, and amber-free Env-tagged N136_TAG_ S401_TAG_, N136_TAG_ S413_TAG_ produced in HEK-293T cells were verified by anti-gp120 western blotting under both suppression and non-suppression experimental conditions. Viral particles of amber-abundant and amber-free HIV-1_Q23 BG505_ were also analyzed using anti-Gag immunoblotting. Viral particles were produced similarly as that used in the infectivity assay, except lack of HIV-1-InGluc. For immunoblot assays, both the supernatants and cells were harvested after 40-48 h post-transfection of HEK293T cells. Virus supernatants were followed by ultracentrifugation of 40,000 rpm for 1.5 h, and virus pellets were resuspended in PBS buffer. Protein samples from virus supernatants and cell lysates were loaded on a 4-12% BisTris polyacrylamide (Bio-Rad, cat#NP0322BOX) gel and transferred onto PVDF membrane by semi-dry blotting (Bio-Rad trans-blot turbo transfer system, 1.4 A constant and 20V for 7 min) with precision plus protein dual color standards (Invitrogen, cat#1610374). PVDF membranes were blocked with 5% (w/v) skimmed milk powder in PBS for 1 h at room temperature and then incubated with primary antibodies of anti-HIV-1-gp120 (NIH HIV Reagent Program, cat #ARP-288), and anti-HIV-1-p24 (NIH HIV Reagent Program, cat#ARP-3537)^85^, diluted in PBST (PBS with 0.05% Tween 20) buffer overnight at 4 °C. Membranes were followed by incubation with HRP-conjugated rabbit anti-sheep/-goat IgG secondary antibodies for 1 h at room temperature. Detection and quantitation of bound antibodies were conducted using clarity western ECL substrate (Bio-Rad, cat#170-5060) and Bio-Rad ChemiDoc™ XRS^+^ System with Image Lab™ Software.

#### Neutralization assays

HEK293T cells were seeded and co-transfected with a mix of plasmids encoding either amber-abundant wildtype or amber-free HIV-1_Q23 BG505_, amber-free N136_TAG_ S401_TAG_, or amber-free N136_TAG_ S413_TAG_, together with HIV-1-InGluc and tRNA^Pyl^/NESPylRS^AF^. 250 µM TCO* was added to the culture medium. At 40 h post-transfection, produced viruses in cell supernatants were collected, filtered, and concentrated by ultracentrifugation. Prior to adding onto the target TZM-bl cells, respective HIV-1 viruses as indicated in figures were incubated with sCD4_D1D2_–lgαtp, PG9 (NIH HIV Reagent Program, cat#ARP- 12149)^58^, PG16 (NIH HIV Reagent Program, cat#ARP-12150)^58^ or PGT151 at the 10-fold serial dilutions (10^2^ to 10^-2^ µg/mL) for 60 min at room temperature, as previously described^33^. Viruses in the presence of different concentrations of ligands were then added on pre-seeded TZM-bl cells in the 48-well plate and were further incubated for another 40-48 h. Approximately two days later, cell supernatants were transferred to 96- well plates for the measurement of Gaussia luciferase activity, as described earlier in the viral infectivity measurements. Relative infectivity (mean ± SD) was normalized to the mean value determined from the control counterpart in each neutralization group.

#### Diameter size measurement of produced virus particles

HEK293T cells were seeded at a density of approximately 2 x 10^6^/well in 10 cm dishes with 12 mL complete growth medium. On the next day, cells with 80% confluency were transfected with 12 µg DNA plasmid and 1 mg/mL PEI Max^TM^ at a molar ratio of 1:3 (w: w). Plasmid DNA and PEI were separately diluted in Opti-MEM reduced serum medium (Thermal Fisher) for 5 min at room temperature (RT). PEI was then mixed well with plasmid DNA and incubated for 15 min at room temperature. The final mixture was added dropwise to the cells. For the amber suppression, HEK293T cells transfected with HIV-1 plasmid DNA, tRNA^Pyl^/NESPylRS^AF^ and PEI at a ratio of 3: 1: 12 (w: w: w) were maintained in complete growth medium supplemented with a final concentration of 250 µM TCO*. The exchange of fresh complete growth medium (supplemented with 250 µM TCO* for amber suppression) was carried out at 6 h post-transfection. At 16 h post-transfection, an additional 250 µM TCO* was added to the amber suppression transfection systems. After 40 h post-transfection, the supernatants containing virions were harvested and centrifugated at a low speed of 200 g for 10 min. The supernatants were purified on a 15% sucrose cushion (w/v phosphate-buffered saline, PBS) by ultracentrifugation at 40,000 rpm for 1.5 h, using Bechman SW-41 Ti Swing Rotor. Virus pellets were resuspended with 200 µL borate buffered saline (BBS) buffer. The diameter size distribution of virus particles diluted in borate buffered saline (BBS) buffer was measured by Nanoparticle Tracking Analysis (NTA) using NanoSight instrument. Three measurements were carried out with 30 seconds duration for each measurement and the data were analyzed by NTA 3.3 Dev Build 3.3.301. Detected tracks were translated into a size distribution using maximum likelihood estimation with an assumed distribution (FTLA method).

#### Preparation of intact HIV-1 virions for smFRET imaging

HIV-1_Q23 BG505_ viruses with Env_BG505_ that is double-tagged (amber-tagged, amber/peptide-tagged, or peptide-tagged) in V1 and V4 loops were prepared for smFRET imaging, in a similar way as previously described^33^. HIV-1 viruses used for smFRET imaging were prepared as replication-incompetent viral particles that lack reverse transcriptase (ΔRT), meaning that all the full-length HIV-1_Q23 BG505_ plasmids used for transfection are ΔRT versions. In this study, we generated three different types of viruses with amber-, amber/peptide-, or peptide-tagged Env: 1) dually peptide-tagged HIV-1_Q23 BG505_ V1Q3 V4A1; 2) dually amber/peptide-tagged HIV-1_Q23 BG505_ V1Q3 S401_TAG_ and HIV-1_Q23 BG505_ N136_TAG_ V4A1 in the amber-free virus context; 3) dually amber-tagged clickable HIV-1 _Q23 BG505_ N136_TAG_ S401_TAG_ and N136_TAG_ S413_TAG_ in the amber-free virus context. The dually peptide-tagged HIV-1 _Q23 BG505_ V1Q3 V4A1 Env carries a Q3 tag in V1 and an A1 tag in V4. As previously described^33, 42^, dually peptide-tagged HIV-1 _Q23 BG505_ V1Q3 V4A1 virions were produced by co-transfecting HEK293T cells with a 40:1 plasmid: plasmid ratio of wild-type HIV-1_Q23 BG505_: Env-tagged V1Q3 V4A1, to ensure that on average only one dually peptide-tagged protomer within a trimer was available for labeling on a single virion. The plasmid: plasmid ratio of untagged: tagged Env was adjusted according to the reduction in amber suppression efficiency of amber-tagged or amber/peptide-tagged Env. Dually amber-tagged Env_BG505_ (N136_TAG_ S401_TAG_ and N136_TAG_ S413_TAG_) or amber/peptide-tagged (V1Q3 S401_TAG_ or N136_TAG_ V4A1) on the amber-free Q23 virions was made similarly, by co-transfecting HEK293T cells with a 4:1 ratio of amber-free HIV- 1_Q23 BG505_ ΔRT to amber- or amber/peptide-tagged ΔRT plasmids, along with an amber suppressor plasmid supplemented with unnatural amino acids. For instance, amber-free HIV-1_Q23 BG505_ N136_TAG_ S413_TAG_ viruses, carrying two clickable sites at N136_TAG_ in V1 and S413_TAG_ in V4, were produced by transfecting an untagged HIV-1_Q23 BG505_ ΔRT genome, an amber-tagged Env variant N136_TAG_ S413_TAG_ ΔRT, and a bi-cistronic plasmid tRNA^Pyl^/NESPylRS^AF^ encoding the amber suppressor tRNA and PylRS^AF^. The amount of tRNA^Pyl^/NESPylRS^AF^ used is 1/3 of the total amount of plasmids that encode Env (including untagged and tagged). The unnatural amino acid TCO* (250 μM) was supplemented to transfected HEK293T cells during the transfection and re-supplemented 12-h post-transfection.

#### Fluorescently labeling Env on intact HIV-1 virions

After 40 h post-transfection, viruses in the supernatants were filtered, harvested, and concentrated on the 15% sucrose cushion by ultracentrifugation at 25,000 rpm. (SW28, Beckman Coulter) for 2 h. Virus pellets were resuspended in the labeling buffer containing 50 mM HEPES, 10 mM MgCl_2_ and 10 mM CaCl_2_^32, 33, 42^. Fluorescently labeling, including enzymatic labeling and click labeling, of virus associated Env has been previously described^32, 33, 42^. For enzymatic labeling, the resuspended HIV-1_Q23 BG505_ V1Q3 V4A1 viruses in labeling buffer were supplied with LD555-cadaverine (0.5 μM, Lumidyne Technologies), LD655-CoA (0.5 μM) (Lumidyne Technologies), transglutaminase (0.65 µM, Sigma Aldrich) ^41^, and AcpS (5 µM)^40^, and then incubated overnight at room temperature. For click labeling, resuspended amber-free HIV-1_Q23 BG505_ N136_TAG_ S401_TAG_ or N136_TAG_ S413_TAG_ viruses that carry two click-chemistry-reactive TCO* were fluorescently labeled in a reaction mixture of 0.1 µM tetrazine-conjugated Cy3 and Cy5 derivatives (LD555-TTZ and LD655-TTZ, Lumidyne Technologies). For enzymatic/click labeling, resuspended amber-free HIV-1_Q23 BG505_ N136_TAG_ V4A1 or V1Q3 S401_TAG_ viruses that carry one peptide and one clickable TCO* were fluorescently labeled by the corresponding enzymatically conjugated dye along with the specific enzyme and the FRET-paired click dye. For the click-labeled or enzymatic/click labeled viruses, a quenching step of 10 min incubation with 1µM BCN-OH quencher followed the labeling step. PEG2000-biotin (Avanti Polar Lipids) was subsequently added to the labeling reaction at a final concentration of 0.1 mg/ml and followed by incubation at room temperature for 30 min with rotation. Excessive label and lipid were removed by ultracentrifugation over a 6–18% Optiprep gradient (Sigma Aldrich) at 40,000 rpm for 1 h. The labeling virions were stored at -80 °C until further study.

#### smFRET imaging data acquisition

All smFRET experiments of Env on intact HIV-1 virions were performed on a customized prism-based total internal reflection fluorescence (prism-TIRF) microscope, as previously described^33, 60, 86, 87^. Fluorescently labeled HIV-1 viruses were incubated in the absence or presence of 0.1 mg/ml sCD4_D1D2_–lgαtp or 0.1 mg/ml PGT151 in the imaging buffer containing 50 mM Tris (pH 7.4), 50 mM NaCl, a cocktail of triplet-state quenchers, 2 mM protocatechuic acid (PCA), and 8 nM protocatechuic-3,4-dioxygenase (PCD) at room temperature for 30 min before smFRET imaging, as previously described^32, 33, 42, 88^. The ligand and antibody concentrations were approximately 5-fold above the 95% inhibitory concentration. HIV-1 viruses carrying fluorescently labeled Env were immobilized on a PEG-passivated biotinylated quartz-coverslip imaging chamber coated with streptavidin. Based on the refractive indexes at the interface between quartz and sample solution, the evanescent field was generated by the total internal reflection of 532-nm single-wavelength laser excitation (Ventus, Laser Quantum) directed to a prism. The donor fluorophore labeled on Env on intact HIV-1 virions is further directly excited by the generated TIRF field. Fluorescence from both donor and acceptor fluorophores labeled on Env was collected through a water-immersion Nikon objective (1.27-NA 60x) and then optically separated by a MultiCam LS image splitter (Cairn Research) with a dichroic filter (Chroma). The separated fluorescence signals from the donor and acceptor passed through two emission filters (ET590/50, ET690/50, Chroma) mounted on the image splitter, respectively. Fluorescence signals were recorded simultaneously on two synchronized sCMOS cameras (Hamamatsu ORCA-Flash4.0 V3) at 25 Hz for 80 seconds. Where indicated, viruses were pre-incubated with the corresponding antibody/ligand for 30 mins at room temperature before imaging and that antibody/ligand was continuously present during smFRET imaging.

#### smFRET data analysis

smFRET data were viewed, processed, and analyzed by a customized SPARTAN software package that was generously shared by Scott Blanchard laboratory (https://www.scottcblanchardlab.com/software)86 and customized MATLAB-based scripts. Image stacks of smFRET data were extracted as individual fluorescence time series trajectories (fluorescence traces) of donor and acceptor labeled on HIV-1 virions. At the single-molecule level, the background signal was evaluated and then subtracted from recorded signals according to the level of signals at the single-step fluorophore bleaching points. Fluorescence traces of the donor and acceptor over 80 seconds were then corrected for the donor to acceptor crosstalk. The time-correlated donor-to-acceptor energy transfer efficiency (FRET values or FRET in graphs) was determined based on FRET= I_A_/(γI_D_+I_A_). I_D_ and I_A_ are the fluorescence of the donor and acceptor, respectively. The correlation coefficient γ compensates for detection efficiencies variations between donor and acceptor channels, as the response of optics and cameras to different wavelengths varies slightly. FRET traces, real-time FRET values over time, were further derived from corresponding fluorescence traces of paired donor and acceptor. FRET traces indicate the relative distance changes between donor and acceptor over time, which are ultimately translated to real-time global conformational dynamics of the donor/acceptor labeled Env in the context of an intact virion. We use stringent criteria to filter out noisy signals. Fluorescence or its corresponding FRET Traces were automatically excluded from further analysis if it lacked signals from either donor or acceptor or both, and if it contained multiple labeled donors/acceptors. After the initial automatic filtering, the remaining traces with sufficient signal-to-noise ratio need to display anti-correlation between donor and acceptor fluorescence before being manually included for further data analysis. The time-correlated anti-correlated feature is a signature of donor-to-acceptor energy transfer in response to real-time conformational changes of the host molecule Env, thus is an indicator of fully active/dynamic Env on the virion. The use of automatic and manual filters ensures that only one dually FRET-labeled protomer within one Env trimer on one virion is included for further analysis. FRET traces that pass our filters were included and compiled into FRET histograms/distributions, reflecting ensembles of multiple conformational states – conformational ensembles. FRET histograms - conformational ensembles represent mean ± SEM, determined from three randomly assigned populations of FRET traces under indicated experimental conditions. FRET histograms were further fitted into the sum of three Gaussian distributions using the least-squares fitting algorithm in MATLAB, based on visual inspection of traces that show clear state-to-state transitions and the idealization of individual traces using 3-state hidden Markov modeling^32, 33, 42, 89^. Each Gaussian distribution represents one distinct conformational state of Env on the virus. The area under each Gaussian estimates the relative probability of Env occupying that conformational state, displayed as relative state occupancy.

**Figure S1.**
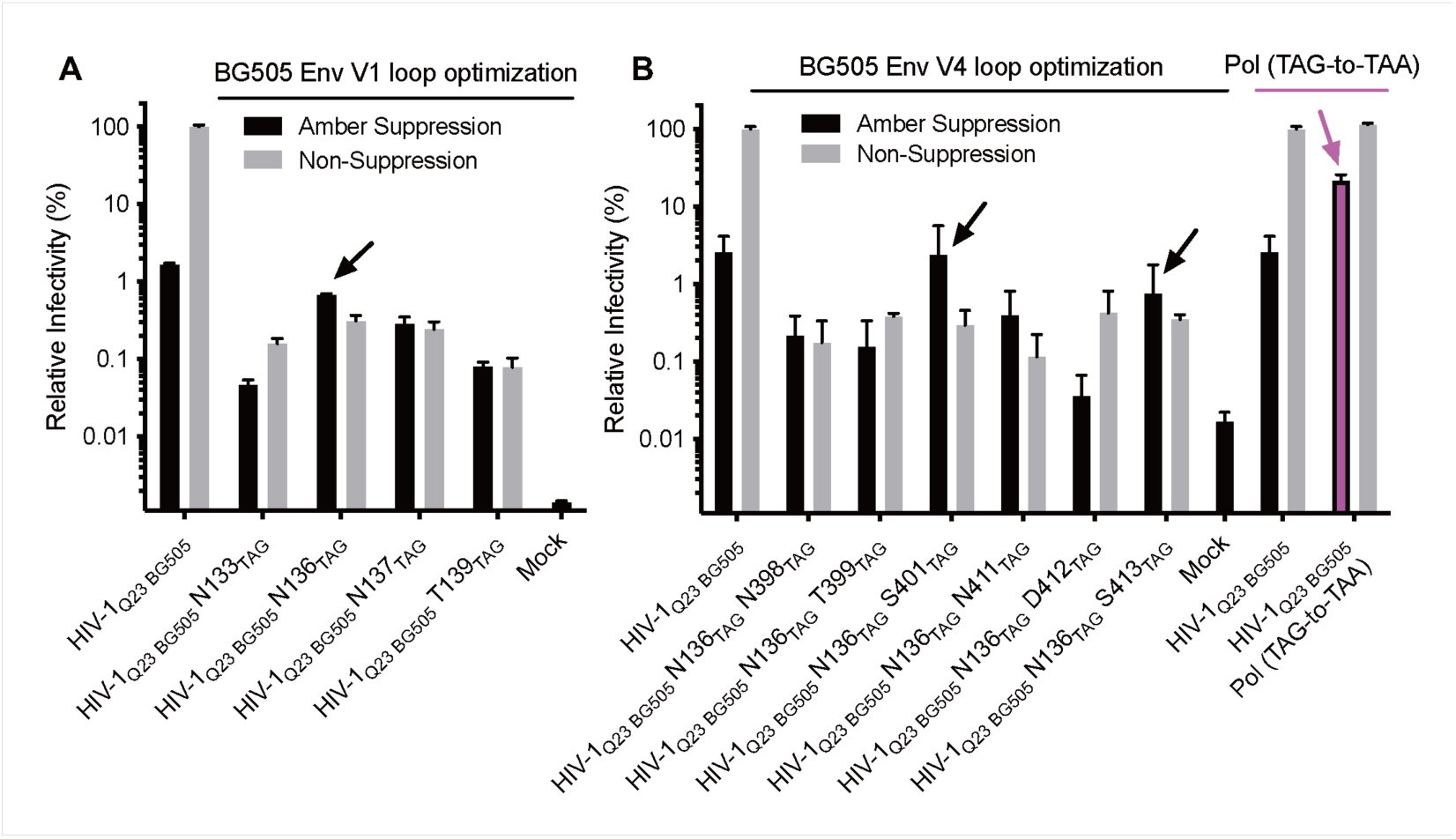
Screening and optimizing amber-tolerable positions in gp120 variable loops V1 and V4 in the naturally existing amber-abundant HIV-1 genomic environment. (A) Infectivity of HIV- 1_Q23 BG505_ carrying single amber termination codon in the genetic sequence of V1 as determined on TZM-bl cells. Infectivity (mean ± SD) was measured from three independent experiments and normalized to that of HIV-1_Q23 BG505_ without amber suppressors/TCO*. The black arrow points to the N136_TAG_, which shows slight suppression activity by amber suppressors. (B) Experiments as in a for single amber insertion in V4 and HIV-1_Q23 BG505_ carrying a termination codon change at Pol from amber to oche (TAG to TAA). Black arrows point to potentially amber-tolerable sites at S401 and S413. The red arrow points to a 5- to 10-fold increase in infectivity caused by the termination codon change at Pol under the suppression condition. This infectivity rescue is a proof of concept of amber issues in the HIV-1 genome shown in Table S2. Results from the HIV-1_Q23 BG505_ (WT) were shown twice in graph (B) for different comparisons.

**Figure S2.**
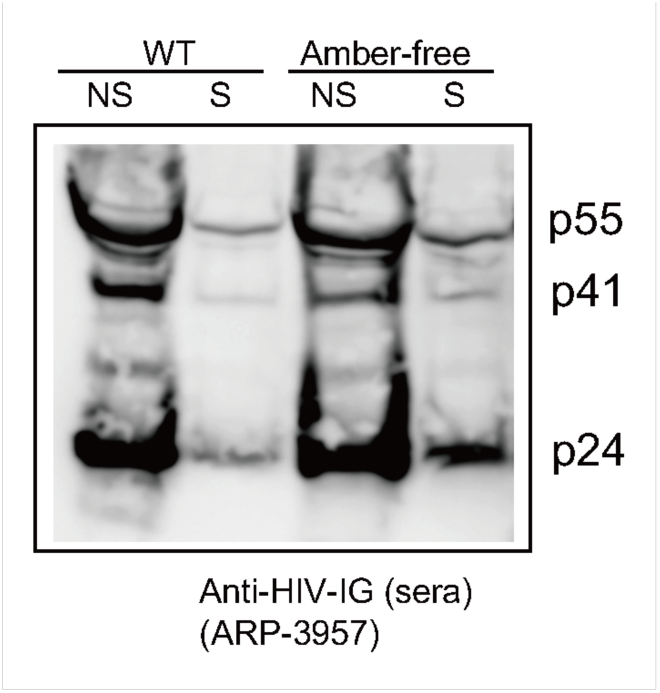
Virus assembly-indicating Gag P24 expression of amber-free and wildtype HIV-1_Q23 BG505_ under different experimental conditions. This analysis was performed by SDS-PAGE of virus particles, followed by western blotting using the indicated polyclonal sera. NS and S stand for the experimental conditions as in Figure 2 of “Non-Suppression” and “Amber-Suppression,” respectively.

**Figure S3.**
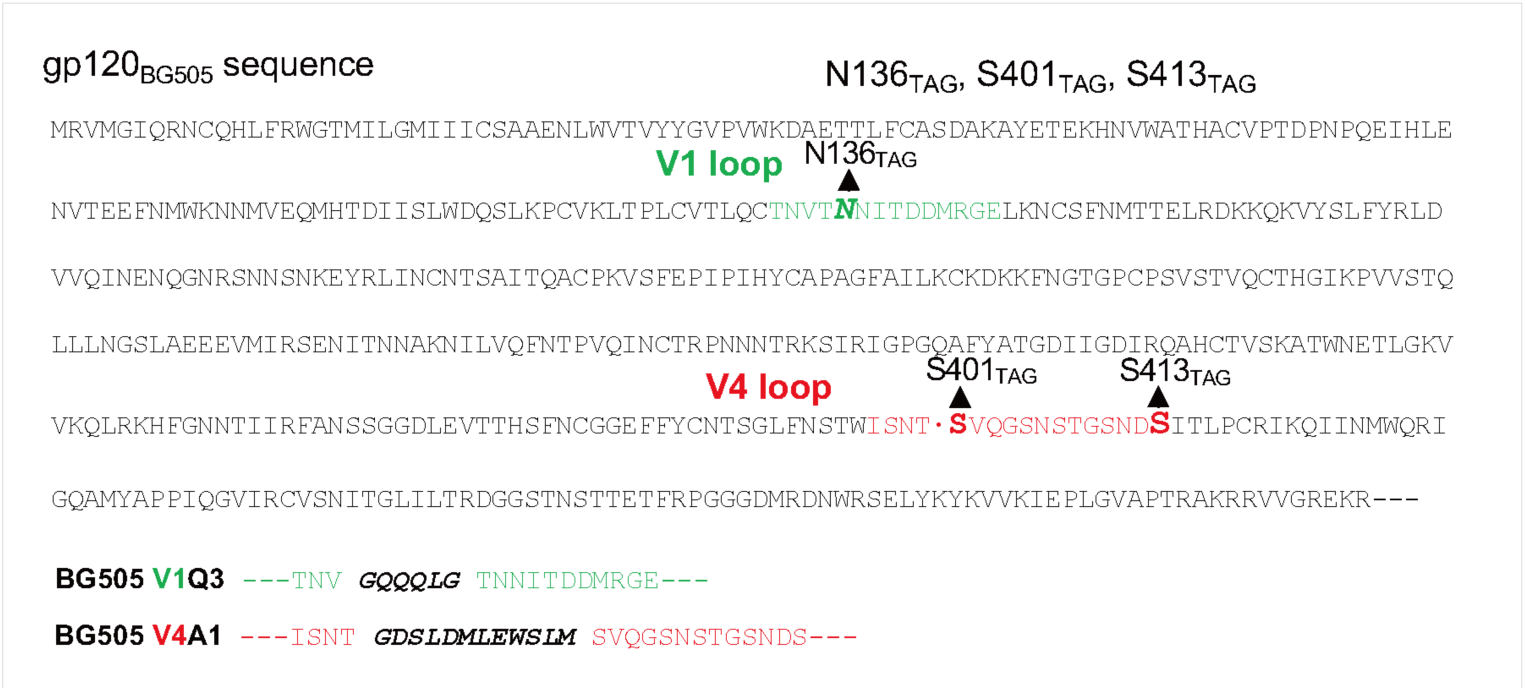
Env_BG505_ gp120 subunit sequence and insertion sites of amber-tag (N136_TAG_, S401_TAG_, and S413_TAG_) and peptide-tag (Q3 and A1) in variable loops V1 and V4.

**Figure S4.**
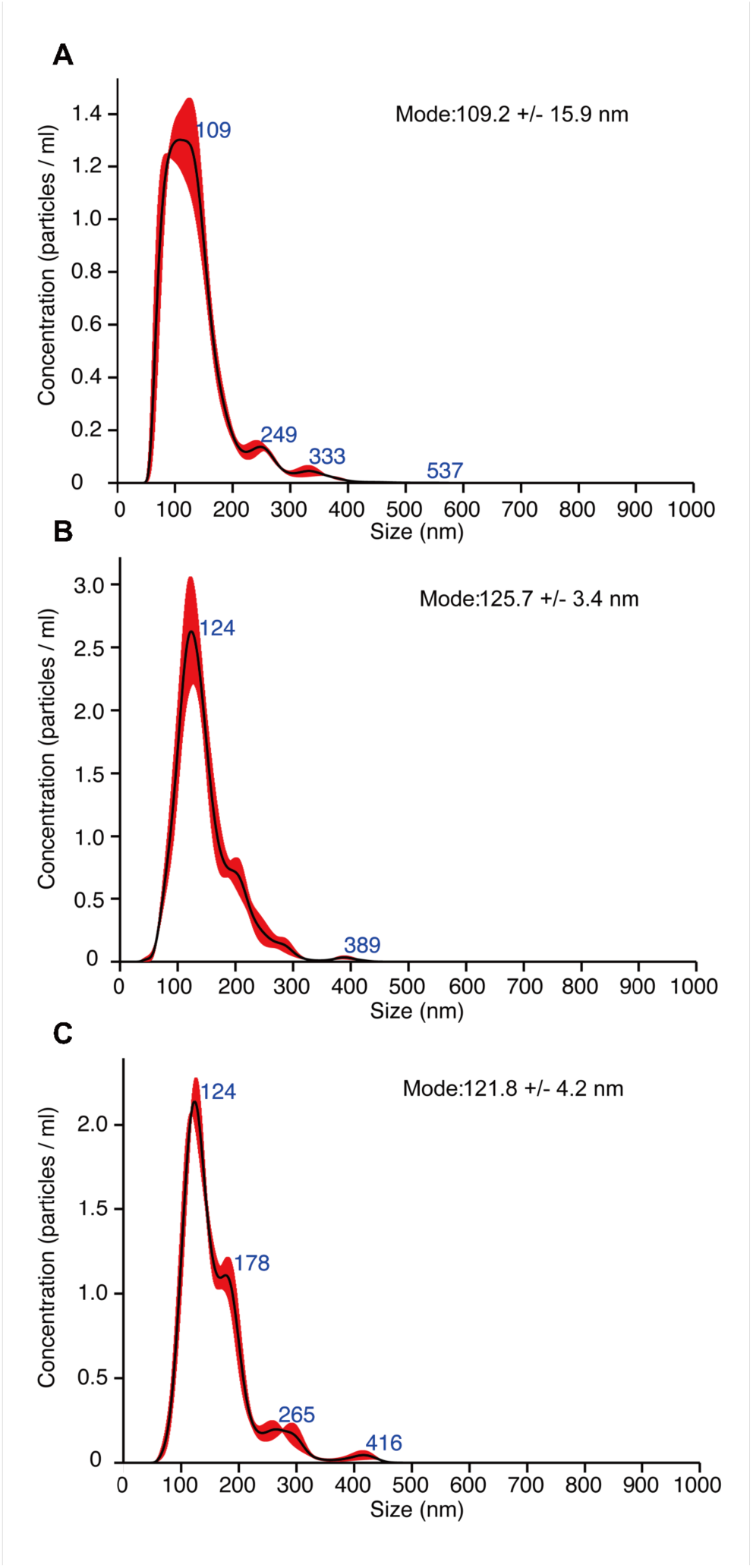
Averaged finite track length adjustment (FTLA) concentration/size of prepared viral particles measured by nanoparticle tracking analysis (NTA). The purified viruses were generated from transfecting HEK293T cells with full-length amber-free HIV-1_Q23 BG505_ (A), and its dually amber-tagged N136_TAG_ S401_TAG_ (B) and N136_TAG_ S413_TAG_ (C) in Env under the suppression condition. Three measurements were conducted with 30 seconds duration and the data were analyzed by NTA 3.3 Dev Build 3.3.301. Detected tracks were translated into a size distribution using maximum likelihood estimation with an assumed distribution (FTLA method). Error bars (+/-) indicate the standard error of the mode size.

**Figure S5.**
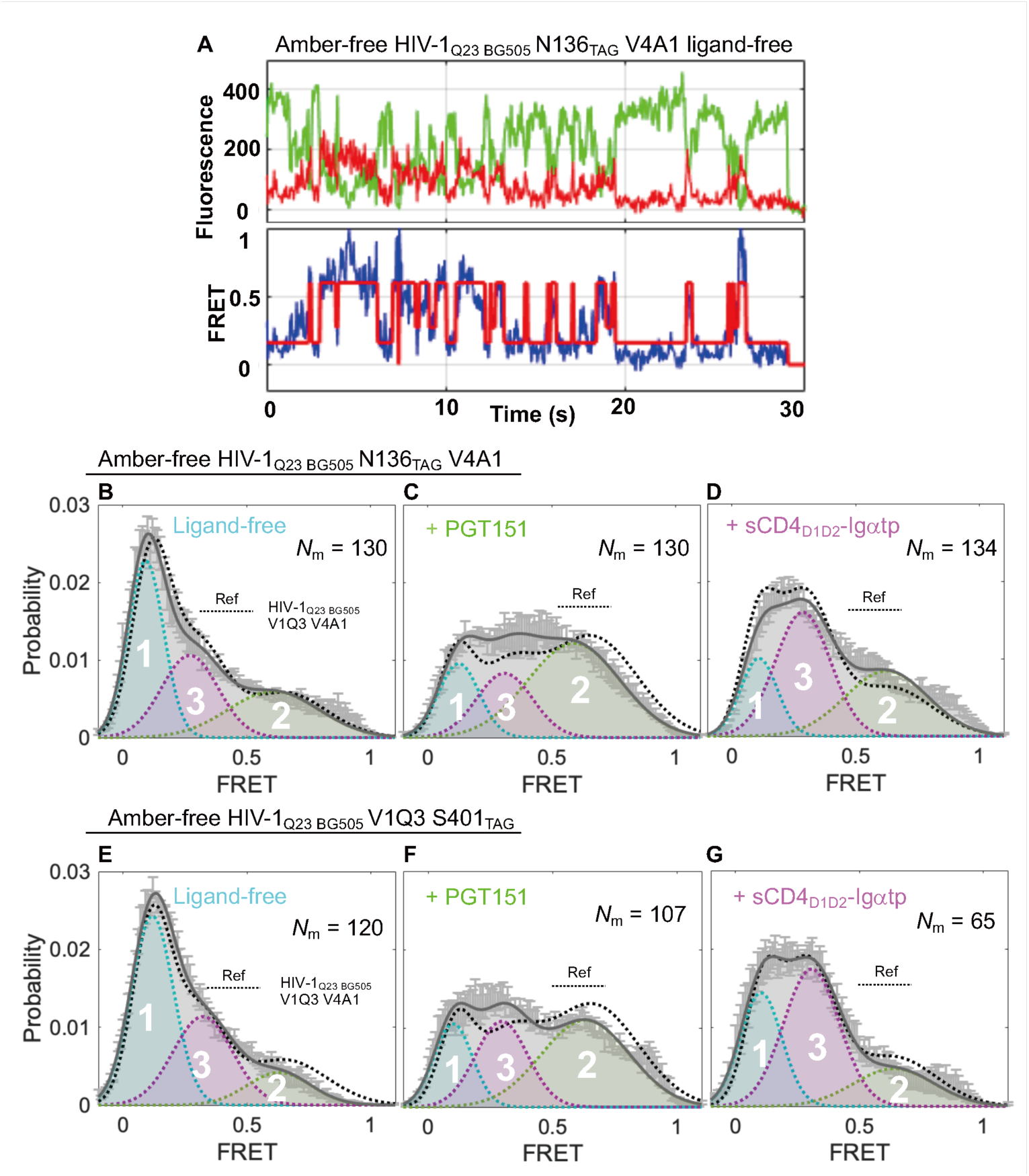
Conformation landscapes of enzymatic/click-labeled Env in the context of amber-free HIV-1 virion resemble those of previously identified based on dually enzymatic-labeled Env. (A) Example fluorescence (Top) and FRET (Bottom) traces for ligand-free enzymatic/click labeled Env_BG505_ on intact virions, HIV-1_Q23 BG505_ N136_TAG_ V4A1. (B–D) Experiments as in Figure 5A–C, respectively, for enzymatic/click-labeled BG505 Env on the amber-free Q23 virion (HIV-1_Q23 BG505_ N136_TAG_ V4A1) in the absence (B) and in the presence of State 3-stabilizer sCD4_D1D2_–Igαtp (C) or State 2-stabilizer PGT151 (D). (E–G) Experiments as in B–D, characterizing the conformational landscapes of ligand-free (E) and upon binding of PGT151 (F) or sCD4_D1D2_–Igαtp (G) to amber-free HIV-_1Q23 BG505_ V1Q3 S401_TAG_. FRET histograms represent mean ± SEM, determined from three randomly assigned populations of FRET traces under corresponding experimental conditions. For evaluated relative occupancy of each conformational state, see Table S3.

**Table S1.**
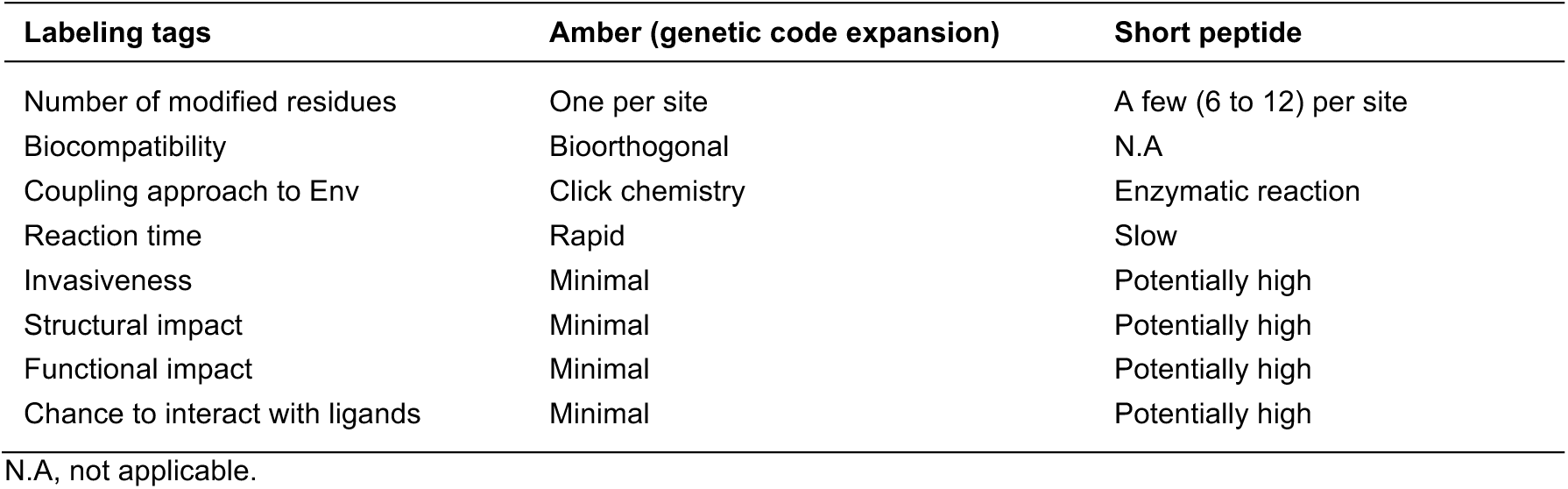
Comparisons between using amber and peptides tags for site-specific labeling of Env

**Table S2.**
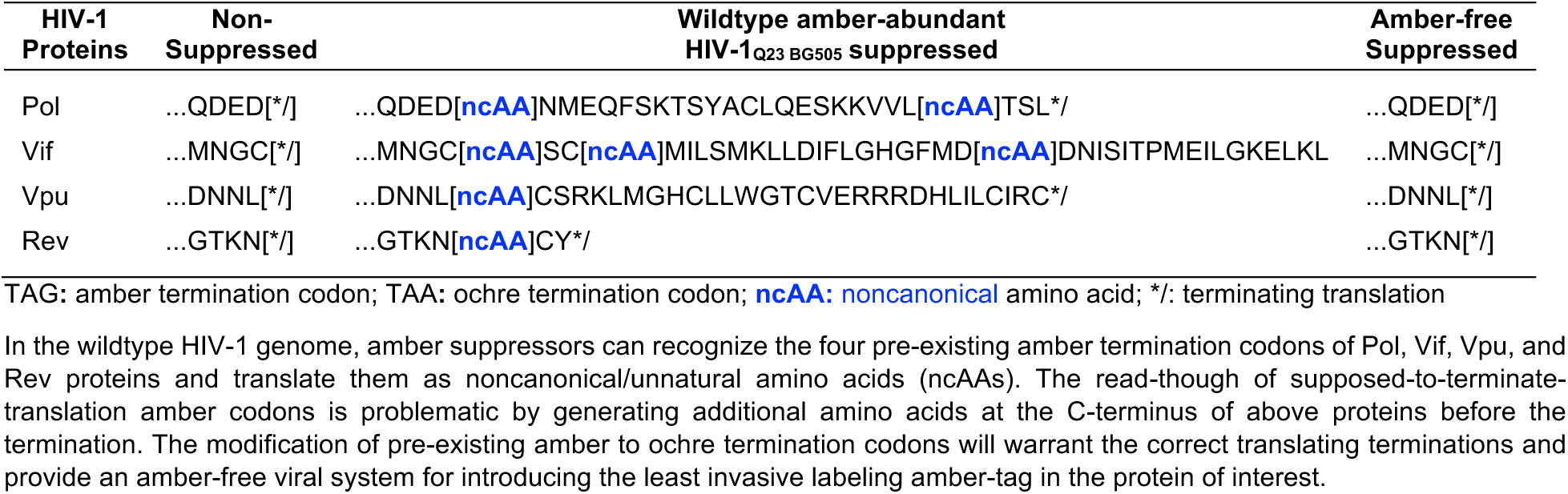
The outcomes of suppressing amber termination codons in wildtype and amber-free HIV-1_Q23 BG505_

**Table S3.**
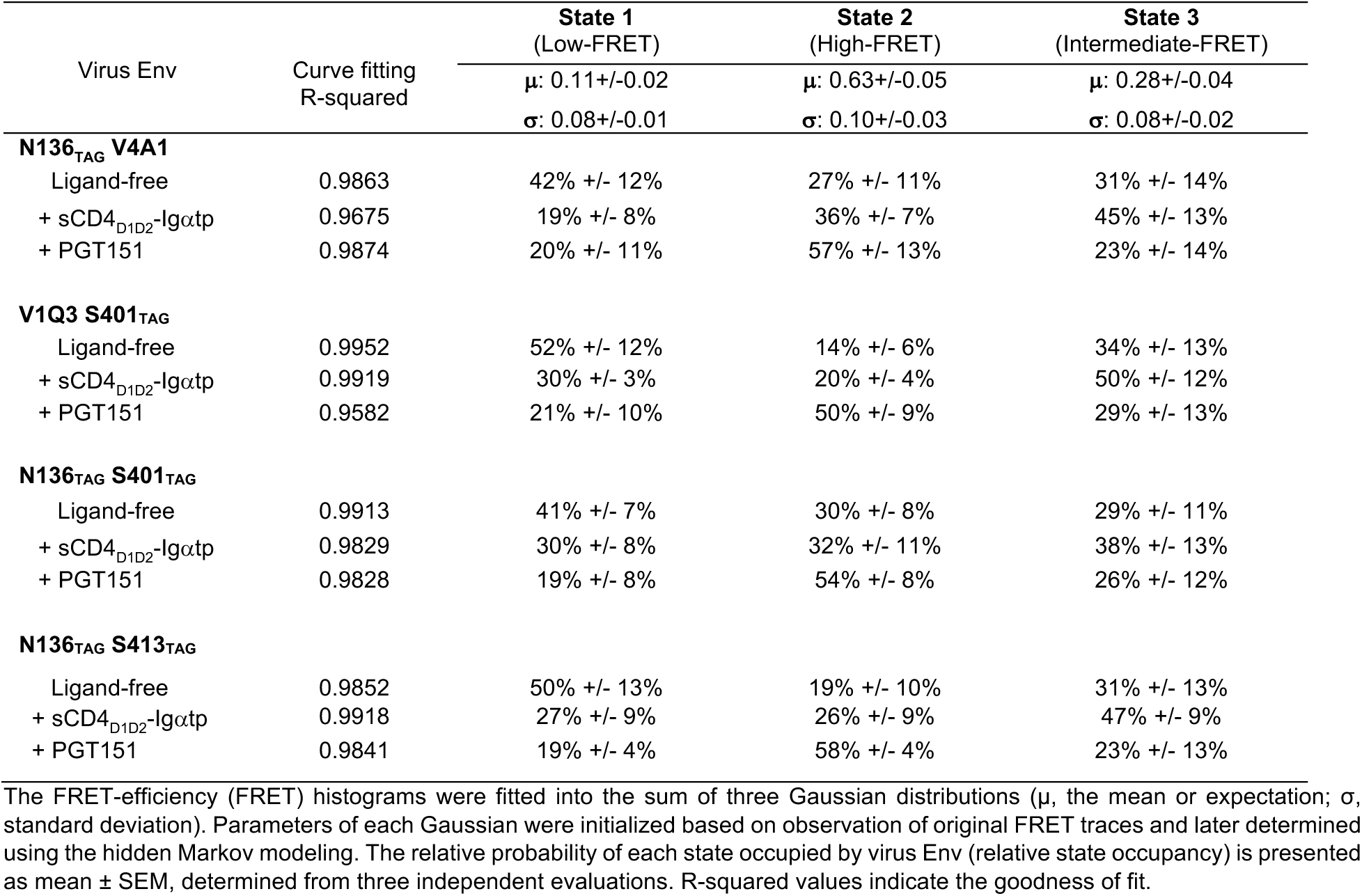
Relative state-occupancy and fitting parameters in each three-state-Gaussian conformational distribution of virus Env

| Virus Env | Curve fitting<br>R-squared | State 1 | State 2 | State 3 |
| --- | --- | --- | --- | --- |
|  |  | (Low-FRET) | (High-FRET) | (Intermediate-FRET) |
| | | $\mu$: 0.11+/-0.02<br>$\sigma$: 0.08+/-0.01 | $\mu$: 0.63+/-0.05<br>$\sigma$: 0.10+/-0.03 | $\mu$: 0.28+/-0.04<br>$\sigma$: 0.08+/-0.02 |
| **N136<sub>TAG</sub> V4A1** |  |  |  |  |
| Ligand-free | 0.9863 | 42% +/- 12% | 27% +/- 11% | 31% +/- 14% |
| + sCD4<sub>D1D2</sub>-Igαtp | 0.9675 | 19% +/- 8% | 36% +/- 7% | 45% +/- 13% |
| + PGT151 | 0.9874 | 20% +/- 11% | 57% +/- 13% | 23% +/- 14% |
| **V1Q3 S401<sub>TAG</sub>** |  |  |  |  |
| Ligand-free | 0.9952 | 52% +/- 12% | 14% +/- 6% | 34% +/- 13% |
| + sCD4<sub>D1D2</sub>-Igαtp | 0.9919 | 30% +/- 3% | 20% +/- 4% | 50% +/- 12% |
| + PGT151 | 0.9582 | 21% +/- 10% | 50% +/- 9% | 29% +/- 13% |
| **N136<sub>TAG</sub> S401<sub>TAG</sub>** |  |  |  |  |
| Ligand-free | 0.9913 | 41% +/- 7% | 30% +/- 8% | 29% +/- 11% |
| + sCD4<sub>D1D2</sub>-Igαtp | 0.9829 | 30% +/- 8% | 32% +/- 11% | 38% +/- 13% |
| + PGT151 | 0.9828 | 19% +/- 8% | 54% +/- 8% | 26% +/- 12% |
| **N136<sub>TAG</sub> S413<sub>TAG</sub>** |  |  |  |  |
| Ligand-free | 0.9852 | 50% +/- 13% | 19% +/- 10% | 31% +/- 13% |
| + sCD4<sub>D1D2</sub>-Igαtp | 0.9918 | 27% +/- 9% | 26% +/- 9% | 47% +/- 9% |
| + PGT151 | 0.9841 | 19% +/- 4% | 58% +/- 4% | 23% +/- 13% |

The FRET-efficiency (FRET) histograms were fitted into the sum of three Gaussian distributions ( $\mu$ , the mean or expectation; $\sigma$ , standard deviation). Parameters of each Gaussian were initialized based on observation of original FRET traces and later determined using the hidden Markov modeling. The relative probability of each state occupied by virus Env (relative state occupancy) is presented as mean ± SEM, determined from three independent evaluations. R-squared values indicate the goodness of fit.

